# On ARGs, pedigrees, and genetic relatedness matrices

**DOI:** 10.1101/2025.03.03.641310

**Authors:** Brieuc Lehmann, Hanbin Lee, Luke Anderson-Trocmé, Jerome Kelleher, Gregor Gorjanc, Peter L. Ralph

## Abstract

Genetic relatedness is a central concept in genetics, underpinning studies of population and quantitative genetics in human, animal, and plant settings. It is typically stored as a genetic relatedness matrix (GRM), whose elements are pairwise relatedness values between individuals. This relatedness has been defined in various contexts based on pedigree, genotype, phylogeny, coalescent times, and, recently, ancestral recombination graph (ARG). ARG-based GRMs have been found to better capture the structure of a population and improve association studies relative to the genotype GRM. However, calculating GRMs and further operations with them is fundamentally challenging due to inherent quadratic time and space complexity. Here, we first discuss the different definitions of relatedness in a unifying context, making use of the additive model of a quantitative trait to provide a definition of “branch relatedness” and the corresponding “branch GRM”. We explore the relationship between branch relatedness and pedigree relatedness through a case study of French-Canadian individuals that have a known pedigree. Through the tree sequence encoding of an ARG, we then derive an efficient algorithm for computing products between the branch GRM and a general vector, without explicitly forming the branch GRM. This algorithm leverages the sparse encoding of genomes with the tree sequence and hence enables large-scale computations with the branch GRM. We demonstrate the power of this algorithm by developing a randomized principal components algorithm for tree sequences that easily scales to millions of genomes. All algorithms are implemented in the open source `tskit` Python package. Taken together, this work consolidates the different notions of relatedness as branch relatedness and by leveraging the tree sequence encoding of an ARG it provides efficient algorithms that enable computations with the branch GRM that scale to mega-scale genomic datasets.

## 1 Introduction

In its most general sense, genetic relatedness refers to the notion of similarity between individuals’ genomes. These similarities are usually summarized as a pairwise comparison of the genomes within an individual and between individuals, or groups of individuals. As a central concepts in genetics, relatedness is used in many applications [Weir et al., 2006, Speed and Balding, 2015]. For example, it has been used to describe genetic variation within and between individuals and groups of individuals in population genetics [Crow and Kimura, 2009, Charlesworth and Charlesworth, 2010], to analyse phenotype covariation between close and distant relatives in quantitative genetics [Falconer and Mackay, 1996, Lynch and Walsh, 1998], and to estimate genetic changes in phenotypic variation over time in evolutionary genetics [Walsh and Lynch, 2018, Arnold, 2023]. For a set of individuals, it is helpful to store their pairwise relatedness in a genetic relatedness matrix, often abbreviated GRM. Over time, genetic relatedness and GRMs have been defined according to pedigree [Fisher, 1919, Wright, 1922], genotype [Cotterman, 1940, Malécot, 1948, 1969, VanRaden, 2008], phylogeny [Felsenstein, 1985, Lynch, 1991], coalescent times [Slatkin, 1991], and recently, ancestral recombination graph [Tsambos, 2022, Fan et al., 2022, Zhang et al., 2023, Tang and Chiang, 2025].

Ancestral recombination graphs (ARGs) describe the network of inheritance relations between a set of individuals via the action of recombination and mutation within a (usually implicit) pedigree [Brandt et al., 2024, Lewanski et al., 2024, Wong et al., 2024, Nielsen et al., 2024], and so provide a common framework in which to consider the various concepts of relatedness. Although ARGs are not directly observable, there has been significant recent progress in inferring ARGs from a sample of DNA sequences [Rasmussen et al., 2014, Speidel et al., 2019, Kelleher et al., 2019, Zhang et al., 2023, Deng et al., 2024, Gunnarsson et al., 2024]. This has been accompanied by computational advances that enable the highly efficient storage and processing of ARGs [Kelleher et al., 2016, Zhu et al., 2024, DeHaas et al., 2024]. In this paper we make use of the *succinct tree sequence* ARG encoding [Ralph et al., 2020, Wong et al., 2024] made available through the `tskit` library.

In addition to providing a unifying framework, ARGs have led to new formulations of relatedness. The “eGRM” of Fan et al. [2022] defines the relatedness between two individuals in terms of the total area of branches in the ARG that are ancestral to both, similar to previous single-tree definitions [Slatkin, 1991]. Fan et al. [2022] showed this is the expected genotype relatedness under a Poisson model of mutation, a special case of a more general duality between “branch” and “site” statistics [Ralph, 2019, Ralph et al., 2020]. The same concept was used by Zhang et al. [2023], although with different terminology, who connected their definition of the “ARG-GRM” to the time to most recent common ancestor (TMRCA) of a single tree [Slatkin, 1991, McVean, 2009]. There are now many different notions of relatedness (see Box 1 for a brief overview), usually defined as an expectation of some quantity (e.g., pedigree relatedness is the expected genetic identity within a pedigree). We therefore use the more precise terms “branch relatedness” and “branch GRM” rather than previously proposed “eGRM” or “ARG-GRM” to avoid confusion.

Recent applications of these methods have highlighted the advantages of using branch information to improve genetic analyses. Fan et al. [2022] demonstrate that the branch GRM (their eGRM) better describes population structure relative to the corresponding genotype GRM, even when based on the same genetic information, and can provide time-resolved characterisations of population structure by considering shared branch areas on particular subsets of the ARG defined by specific time intervals. Link et al. [2023] applied a branch GRM to improve mapping of quantitative trait loci in the presence of allelic heterogeneity and in understudied populations. Tang and Chiang [2025] modified the “eGRM” to better reveal the recent relatedness among admixed individuals. Tsambos [2022] developed a method to find DNA segments that are identical-by-descent (IBD) for pairs of individuals in a given ARG and then summarise these outputs, possibly as an “IBD GRM”, which provides an ARG-based analogue to the pedigree GRM. Zhang et al. [2023] use a branch GRM (their ARG-GRM) to estimate heritability and to perform a “genealogy-wide association scan”, showing that this approach can be more powerful at detecting the effect of rare variants than association analysis on SNP array genotypes imputed to whole-genome sequence genotypes. Gunnarsson et al. [2024] extended this work to a large whole-genome sequence dataset and Zhu et al. [2024] used randomized linear algebra to scale the estimation of heritability and region-based association testing with branch GRM.

The scalability of current exact ARG-based relatedness methods, however, is constrained by their need to generate and store the full branch GRM. As the GRM encodes all pairwise relationships among *n* samples, it requires at least *O*(*n*^2^) time and space to compute. Several currently available datasets of core interest for these methods consist of hundreds of thousands of samples [Caulfield et al., 2017, Turnbull et al., 2018a, Bycroft et al., 2018, Backman et al., 2021, Ros-Freixedes et al., 2022b, Halldorsson et al., 2022, UK Biobank Whole-Genome Sequencing Consortium et al., 2023, All of Us Research Program Genomics Investigators et al., 2024], and genomic datasets with millions of samples are increasingly available [e.g. Cesarani et al., 2022, Stark et al., 2024, Cook et al., 2025, Cole et al., 2025]. At this scale, algorithms with quadratic time and space complexity are simply not feasible. However, the GRM itself is often not the goal; rather, we are usually interested in what we can do *with* the GRM. For example, population genetic applications such as principal component analysis (PCA) and quantitative genetic applications such as estimation of heritability, are defined in terms of core linear algebra operations performed with the GRM, and the outputs are of much smaller dimension. Given that all the information in a GRM is encoded in an ARG, there is the possibility that we can bypass generating large intermediate matrices and instead compute the quantities of interest directly. This approach was used by Zhu et al. [2024], who use the ARG for fast, approximate GRM-vector multiplication. Indeed, the ARG can be seen as a sparse matrix representation of the genotype matrix which can hence naturally be used for efficient computation [Ralph et al., 2020].

In this paper, we begin by defining a trait-centric concept of genetic relatedness, following long-standing approaches in the field [Fisher, 1919, Wright, 1922]. We show how branch relatedness arises as the covariance of a trait determined additively along the branches of an ARG, and how this relates to other measures of relatedness. We then illustrate these definitions and the relationships between pedigree relatedness and branch relatedness using simulated data from a real pedigree of French-Canadian individuals. Next, we describe a relatively efficient algorithm to compute the entire branch GRM that has complexity *O*(*tn*^2^), where *t* is the number of local trees (or equivalently, number of recombination breakpoints) in an ARG. As discussed in the previous paragraph, explicit representations of the entire GRM are necessarily limited in scale, so we turn to matrix-vector products. We then present an algorithm to compute the product of the branch GRM with an arbitrary vector, and show that it has *O*(*n* + *t* log *n*) time complexity and *O*(*n*) space complexity. We can therefore compute branch GRM-vector products substantially faster and with less memory than the branch GRM itself. We illustrate the utility of this approach by presenting a randomized singular value decomposition method for PCA of the branch GRM (implemented in `tskit`), and show that it scales to millions of samples via benchmarks.

## 2 Results

### 2.1 ARGs and tree sequences

We first introduce our notation for ARGs, following Kelleher et al. [2016, 2018], and Wong et al. [2024]. Using this terminology, an ARG represents the history of a set of sampled genomes by a collection of *nodes* and *edges*. Each chromosome of an individual is represented by a *node*, and each node has an associated *time*, indicating when the individual was born. In diploids, the two haploid genomes of a genotyped individual are represented by two *sample* nodes. *Ancestral* nodes represent genomes of non-genotyped individuals. Each edge encodes the inheritance of some genome segment by some “child” node from a “parent” node (despite the terminology, these two may be separated by more than one generation). Edges are also often referred to as *branches*. The *span* of an edge is the length of the inherited segment of genome, and the *length* of an edge is the number of generations across which the segment was inherited, that is, the difference between the times of parent and child nodes. Genetic variation is represented in this structure by recording where in the ARG mutations occurred. For example, if we say that a mutation that produces a C nucleotide occurs at genomic position *x* on the edge from some parent *p* to a child *c*, then (1) the mutation has occurred somewhere in the chain of inheritances by which *c* has inherited the genetic material from *p*; and (2) any other nodes that inherit from *c* at position *x* will carry a C, unless another mutation intervenes. Finally, recombination events are implicitly encoded by edges between child and parent nodes, that is, a child node can inherit from different nodes of different parents. Inheritance relationships at each location of the genome are described by a *local tree*, and subsequent local trees are separated by the genomic locations of recombination events.

The *succinct tree sequence*, or *tree sequence* for short, is an efficient ARG encoding [Kelleher et al., 2016, 2018, Wong et al., 2024]. The data structure is based on a succinct description of nodes, edges, and mutations as described above, and can be used to efficiently recover and process the sequence of local trees that describe how the samples are related on each consecutive section of the chromosome.

### 2.2 A trait-centric notion of genetic relatedness

Consider an additive trait, that is, a trait whose value is the sum of effects associated with each allele carried by the individual. Suppose that the genotypes at each locus are from some alphabet *A*, and that at each locus ℒ in the genome there is an “ancestral” allele *a*_ℒ_. The additive effect of allele *x* at locus ℒ is *Z*_ℒ,*x*_, which is relative to the ancestral allele, so that 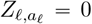. More discussion of this choice can be found in Appendix A. Then, an individual’s genetic value is the sum of the effects of alleles across all *n*_*L*_ loci, averaged across genome copies. We will write *G*_*i*,ℒ,*g*_ for the allele of the *g*^th^ genome copy of individual *i* at locus ℒ, so that a *p*-ploid individual *i* has genetic value:

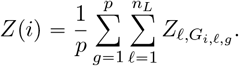

Finally, suppose that the effects of each non-ancestral allele *Z*_ℒ,*x*_ are independently drawn from a probability distribution with mean zero and variance *σ*^2^. The choice to average across genome copies (as opposed to, say, sum them) is only consequential for situations where mixed ploidy is considered and implies a particular model of dosage compensation or how we summarise relatedness across genome copies. Mixed ploidy arises with sex chromosomes or haplodiploids [Grossman and Eisen, 1989], or summarizing relatedness between groups with different number of individuals [Cockerham, 1967]. Many measures of relatedness make use of this trait model (explictly or implictly), in which case relatedness is proportional to covariance between individuals’ trait values. We now demonstrate this equivalence.

For simplicity, suppose for the moment all loci are bi-allelic, so *G*_*i*, ℒ,*g*_ ∈ {0,1}, and *Z*_ℒ,0_ = 0. See Appendix B for a more general discussion. Under this model, if we write *p*(*i*, ℒ) as the proportion of alleles carried by individual *i* at locus ℒ that are not ancestral (so *p*(*i*, ℒ) = (*G*_*i*,ℒ,1_ + *G*_*i*,ℒ,2_) / 2 for diploids), then the covariance between the traits of individuals *i* and *j* is:

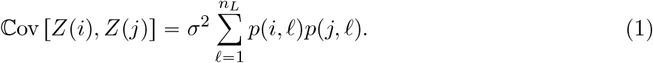

Note that this is the covariance of *Z*(*i*) and *Z*(*j*) as random variables, averaging over random assignment of allelic effects, but with genotypes fixed (*p*(*i*, ℒ) is not random).

The above covariance expression (1) depends on the choice of ancestral allele. This seems undesirable for a measure of relatedness; choosing a point farther back in time as a reference, so that a different allele is “ancestral” and the derived allele is likely fixed, should not affect relatedness within the population. It does affect the relatedness calculated above because this is, implicitly, a model of trait variation *relative* to a hypothetical individual whose genotype is composed entirely of ancestral alleles. A common approach to resolve this is to *center* the traits, which takes the mean of some individuals as reference, rather than a hypothetical ancestor. When using pedigree data, these reference individuals are founders of the pedigree [Wright, 1922], while when using genotype data these reference individuals are the genotyped individuals [VanRaden, 2008].

Suppose we have *n*_*I*_ haploid individuals 1, …, *n*_*I*_ with genetic values *Z*(1), …, *Z*(*n*_*I*_), and define the mean allele frequency among these individuals as 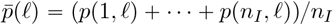. The covariance of the traits after centering to the sample mean is:

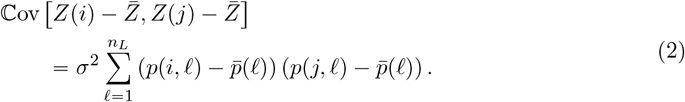

If the derived allele at locus ℒ is fixed, then 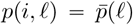 for all *i*, and so such loci do not contribute to the covariance expression (2).

It will be helpful to use another form of the mean centering of expression (2). If *U* and *V* are random, uniformly chosen individuals from the sample, and *L* a random, uniformly chosen locus, then, we can rewrite 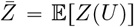 and 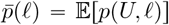. Consequently, the two sides of (2) are also equal to:

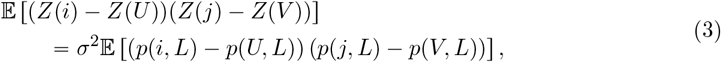

where the expectation is averaging over choice of *U, V*, and *L*.

The expression (3) highlights a connection to the familiar genotype GRM. Simplifying to haploids, we can treat 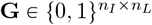 as the genotype matrix for *n*_*I*_ haploid individuals at *n*_*L*_ loci. We are interested in the covariance between individuals *i* and *j*, that is, between the two genomes in rows *i* and *j* of **G**. Let **G**^*c*^ be the *column-centered* haplotype matrix with entries 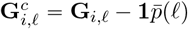. A common definition of covariance is:

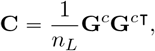

so that the covariance between individuals *i* and *j* based on their genotypes is:

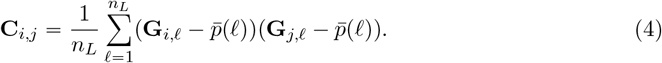

This expression (4) is the kernel of many variants of GRM [VanRaden, 2008, Yang et al., 2010, Speed and Balding, 2015, Zhang et al., 2023], apart from difference between the haploid and diploid setting, with the latter being an aggregate form of the former [Cockerham, 1967, Smith and Allaire, 1985]. This expression (4) is also equal to (2) divided by *n*_*L*_, after setting *σ*^2^ = 1. The corresponding expression for diploids uses in place of **G** the allelic dosage matrix whose entries are the proportion of non-reference alleles carried by the individual. It is more common in the literature to define the allelic dosage matrix as the *number* of non-reference alleles; here we define it as the proportion so that it agrees with (2); this is necessary because of the convention to define *Z*(*i*) as the average across the *p* genome copies. For diploids this results in an additional factor of four.

Many definitions of relatedness weight the contribution of the ℒ^th^ locus by 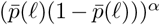. We take *α* = 0 for simplicity, but the discussion below applies more generally.

A third interpretation of this covariance can be derived as follows. As before, take *i* and *j* are two fixed haploid individuals; also take two additional random haploid individuals *U* and *V* and form the random variable (*X*_*i*_, *X*_*j*_, *X*_*U*_, *X*_*V*_) that takes the value (*G*_*i*,ℒ_, *G*_*j*,ℒ_, *G*_*U*,ℒ_, *G*_*V*,ℒ_) with probability 1 / (*n*_*I*_ ^2^*n*_*L*_) for *U, V* = 1, …, *n*_*I*_ and ℒ = 1, …, *n*_*L*_. In other words, we choose the individuals *U* and *V* uniformly at random, with replacement, from the set of *n*_*I*_ individuals, and also choose a locus ℒ uniformly at random from the set of *n*_*L*_ loci; then (*X*_*i*_, *X*_*j*_, *X*_*U*_, *X*_*V*_) is the alleles of those individuals at that locus. Then, as shown in Appendix C, it turns out that

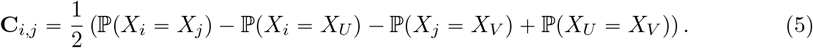

This expression is more readily extendable to multi-allelic data.

We therefore have the following three equivalences (2, 4, and 5):

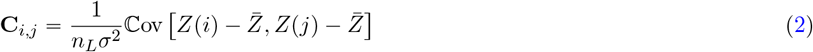

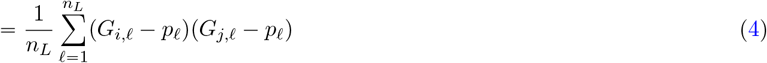

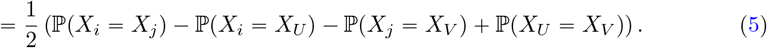

From the third equivalence (5), the quantity *n*_*L*_**C**_*i,j*_ has the following interpretation. Let *m*(*i, j*) denote the number of pairwise allele matches between the individual *i* and *j*, and let *U* and *V* be independently chosen individuals from the set of individuals. Then the quantity *n*_*L*_**C**_*i,j*_ is the expected number of pairwise allele matches between *i* and *j* relative to the rest of individuals:

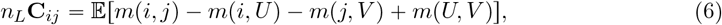

where the expectation is over the choice of U and V. This interpretation is closely related to the definition of kinship between individuals *i* and *j* as the “the probability of a match between alleles drawn at random from each of them”, averaged over loci, and with the alleles drawn with replacement if *i* =*j* [Malécot, 1948, *1969, Speed and Balding, 2015]*. *See also Weir and Goudet [2017, 2018] and Ochoa and Storey [2021] on other “relative” kinship estimators*.

### 2.3 A trait-centric perspective on the branch relatedness

We now describe a closely related notion of branch relatedness. Suppose that we only observe the relationships in the ARG, not the mutations that appear in it. This is similar to the starting point of pedigree relatedness, but we assume we also know full ancestry of each genome all the way to the roots of each local tree (the MRCAs) and which portions of the genomes were inherited in each relationship. The expected number of mutations that appear on a segment of genome of *s* base pairs inherited across *b* generations is proportional to *b* × *s*. In other words, the expected number of mutations on an edge of length *b* and span *s* is proportional to its area, *A* = *b* × *s*. If the effect of each mutation has variance *σ*^2^, then the variance of the edge effect is *Aσ*^2^. (This is because the variance of the sum of a random number *N* of independent and identically distributed mean-zero terms is the mean of *N* multiplied by the variance of the terms.) Let *A*(*i, j*) be the total area of branches ancestral to individuals *i* and *j*. Then, just as above, with randomly chosen individuals *U* and *V*:

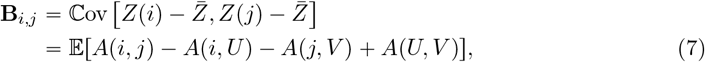

where the expectation in the second line is over the choice of *U* and *V*, while *Z*(*i*) and 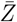 are defined as before. Note that the random variables *Z*(*i*) are the same as before, but this expression differs from (2) in that here the covariance averages not only over allelic effects, but also over location of the mutations. This is an example of a general relationship described in Ralph et al. [2020].

The branch relatedness can also be rewritten as a weighted average of coalescence times, as noted by McVean [2009], Fan et al. [2022], and Zhang et al. [2023]. Let *s*_*k*_ be the genome sequence length corresponding to the *k*^th^ local tree, and within this tree define *b*(*i, j, k*) be the total length of branches ancestral to both haploid individuals *i* and *j, t*(*i, j, k*) be the TMRCA of *i* and *j*, and 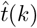 the time of the root. Supposing that *i* and *j* are both at time 0, then the time of the root is equal to the TMRCA plus any additional, shared, branch lengths:

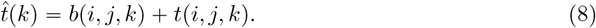

We can use this relationship to split (7) by local tree as follows:

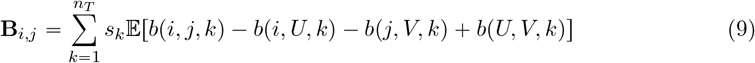

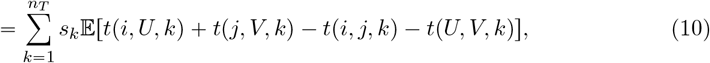

where *n*_*T*_ is the number of local trees in the ARG, and the expectation averages over *U* and *V*.

Our definition differs slightly from the eGRM by Fan et al. [2022] Let *S*_*i,e,k*_ = 1 if sample *I* is a descendant of branch *e* in the *k*^th^ tree and *S*_*i,e,k*_ = 0 otherwise, and 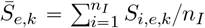 for the proportion of samples inheriting from *e*. Also, write *b*_*e*_ for the length of *e*. Then (7) can be rewritten as:

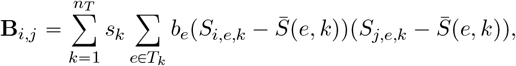

where the second sum is over edges *e* in the *k*^th^ tree. On the other hand, Fan et al. [2022] define the eGRM as:

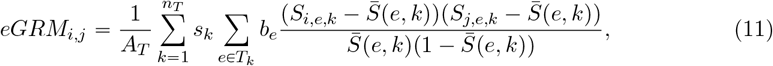

where 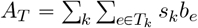 is the total area of the ARG. The different denominator normalises the contribution of each edge according to its standard deviation in the population, and is equivalent to the 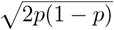 standardization often used in the genotype GRM.

Relatedness and divergence are closely related, as demonstrated by the relationship (8). Let *d* (*i, j, k*) be the distance in the *k*^th^ tree between *i* and *j*, and *r*_*k*_ the root of the *k*^th^ tree. A more general relationship that does not assume *i* and *j* are both at time zero is [Semple and Steel, 2003]:

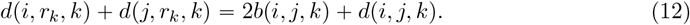

That is, the sum of the distances from each to the root is equal to the distance between them plus twice the distance from their MRCA to the root. Let *R*(*i*) denote the sum along the genome of the distances from *i* to the root and *D*(*i, j*) the sum along the genome of the distances between *i* and *j* in the local trees. Then *D*(*i, j*) is the branch genetic divergence between *i* and *j* [Ralph et al., 2020], and summing the previous relation across the genome, we get:

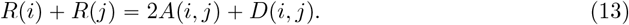

Rearranging and substituting into the expression for branch relatedness (7), centering cancels the terms with *R*, giving:

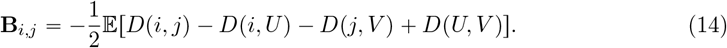

Thought of as matrices, if **P** = **I** − **11**^**⊤**^ /*n*_*I*_ is the *n*_*I*_ × *n*_*I*_ centering matrix, the above equation says that **B** = − **PDP**/2. For more discussion and other relationships between relatedness and divergence, see Zhang et al. [2023], Supplementary Note 3.

### 2.4 Branch PCA

Principal component analysis (PCA) is a commonly used technique to quantify and visualise population structure from genotype data. Mathematically, PCA operates by projecting samples onto a set of orthogonal axes, each defined as a linear combination of genotype values across SNPs or other genetic variants. An iterative characterization of PCA is as follows: choose the first principal component to be the axis that captures the maximum possible variance in the data, then choose the second principal component that maximizes variance whilst being orthogonal to the first, and so on. The first three or four principal components are often presented as a low-dimensional summary of population structure.

Principal components can be found, equivalently, as eigenvectors of the (centered) genotype GRM or singular vectors of the underlying (centered) genotype matrix. Both decompositions can be efficiently approximated with randomized algorithms that can operate on matrices only implicitly defined through matrix-vector products [Halko et al., 2011]. McVean [2009] gave a genealogical interpretation of PCA, while Fan et al. [2022] showed that branch PCA can in some cases better capture recent population structure than genotype PCA, even when based on the same genotype information. In sub-section 2.7 we describe an efficient algorithm for such a product that bypasses the construction of branch GRM and operates directly on tree sequence encoding of an ARG. However, we first discuss the connection between pedigree and branch relatedness.

### 2.5 Connection between pedigree and branch relatedness

The pedigree relatedness of two individuals in a given pedigree is the probability that they both inherit from the same ancestral genomes within the pedigree at a given locus, that is, that they are IBD within the pedigree [Malécot, 1969]. While pedigree and branch relatedness seem similar, they in fact differ in what they measure. Using IBD to define an ARG-based notion of relatedness would lead to an “IBD GRM” in the sense discussed by Tsambos [2022]. While the IBD relatedness measures the probability of identity with reference to a particular time (or set of ancestors) in the past, branch relatedness measures shared edge area. This difference in units is confusing given the correspondence between the expressions (5) and (7). Because branch relatedness **C**_*i,j*_ can be written using probabilities of identity, it seems analogous to IBD relatedness within a pedigree or an ARG, but the close theoretical relationship to **B**_*i,j*_ tells us that it is better to think about branch relatedness as having units of “shared time”.

### 2.6 Demonstration with the French-Canadian pedigree

To empirically illustrate the connection between pedigree and branch GRM, we analysed the pedigree of a subset of 2,321 individuals from the BALSAC dataset, drawn from five different regions in Quebec (Figure 1G). For this subset, we computed pedigree and branch GRM. For the latter, we obtained an ARG from pedigree- and ancestry-informed simulation and computed branch GRM from the ARG. We simulated 100 such ARGs to evaluate the variance in branch GRM within a fixed pedigree. See Methods for more details on the dataset and simulations performed.

**Figure 1:**
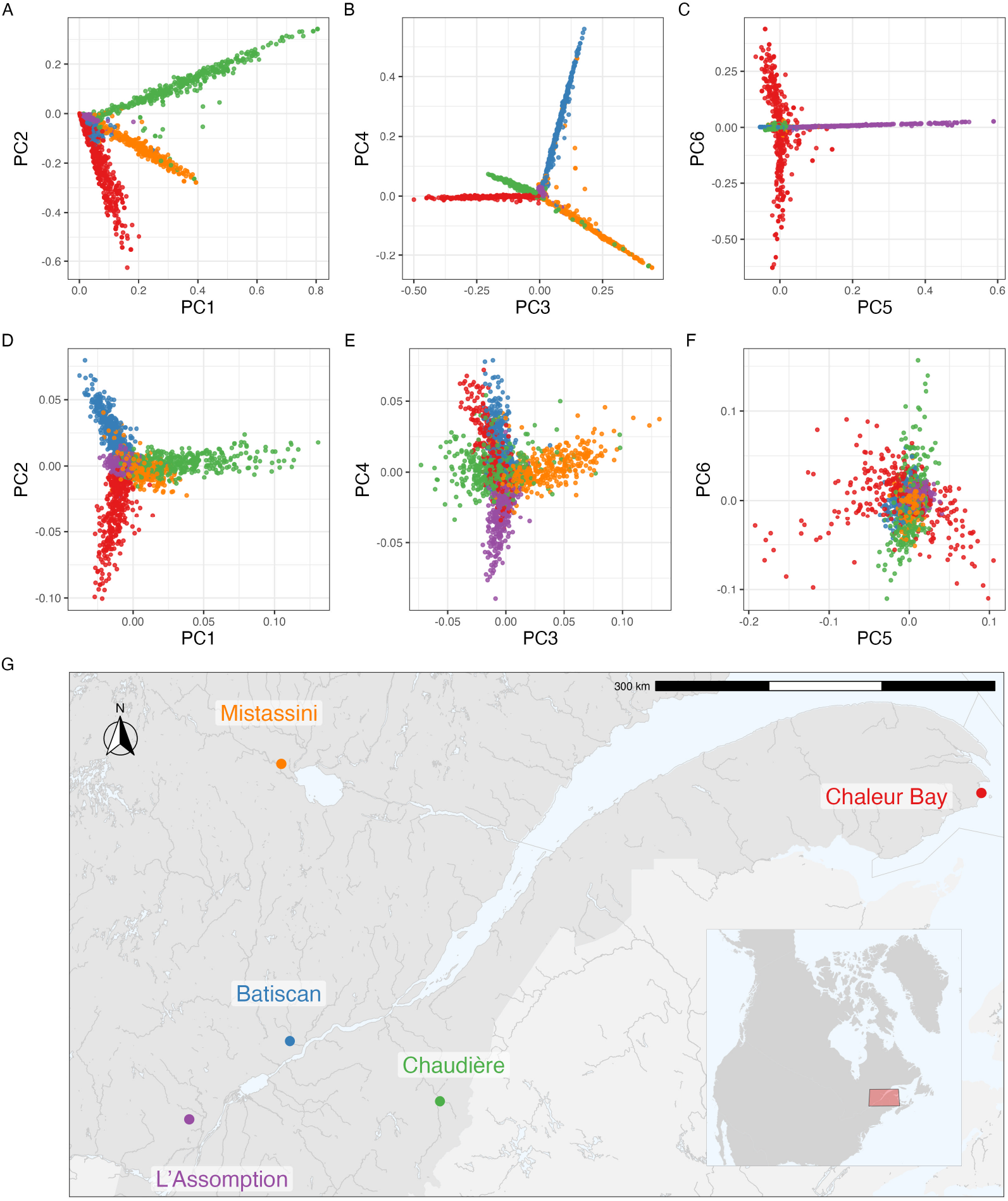
Principal components analysis of pedigree and branch GRM of French-Canadian individuals. (A-C) The first six principal components of pedigree GRM. (D-F) The first six principal components of branch GRM. (G) A partial map of Quebec with approximate locations of sampled individuals.

The overall population structure according to pedigree and branch PCA of the 2,321 individuals is shown in Figure 1 with noticeable clustering by the five regions. The pedigree PCs show sharp distinctions between individuals from different regions (Figure 1A-C). Although all regions share a common bottleneck, over the last four centuries there has been a sufficiently little movement that each region pulls a distinct principal component. PCs 1 and 2 show a clear structure between Chaudière, Chaleur Bay, and Mistassini, PCs 3 and 4 distinguish between each of the five regions except L’Assomption, and PCs 5 and 6 illustrate a clear distinction between L’Assomption and Chaleur Bay.

The branch PCs also show the population structure (Figure 1D-F), but with substantially less “clean” patterns. This higher variation is due to the randomness of genetic inheritance within a single chromosome, which is averaged over by pedigree relatedness [Weir et al., 2006, Hill and Weir, 2011, Thompson, 2013, García-Cortés et al., 2013]. This variability is also evidenced when comparing relatedness between a subset of 250 individuals: a clearer structure emerges with pedigree GRM than with branch GRM (Figure 2). Notably, when individuals have a shallow pedigree (for example, one sampled individual from Chaleur Bay), their corresponding pedigree GRM rows and columns have low pedigree relatedness, while this is not the case with branch relatedness.

**Figure 2:**
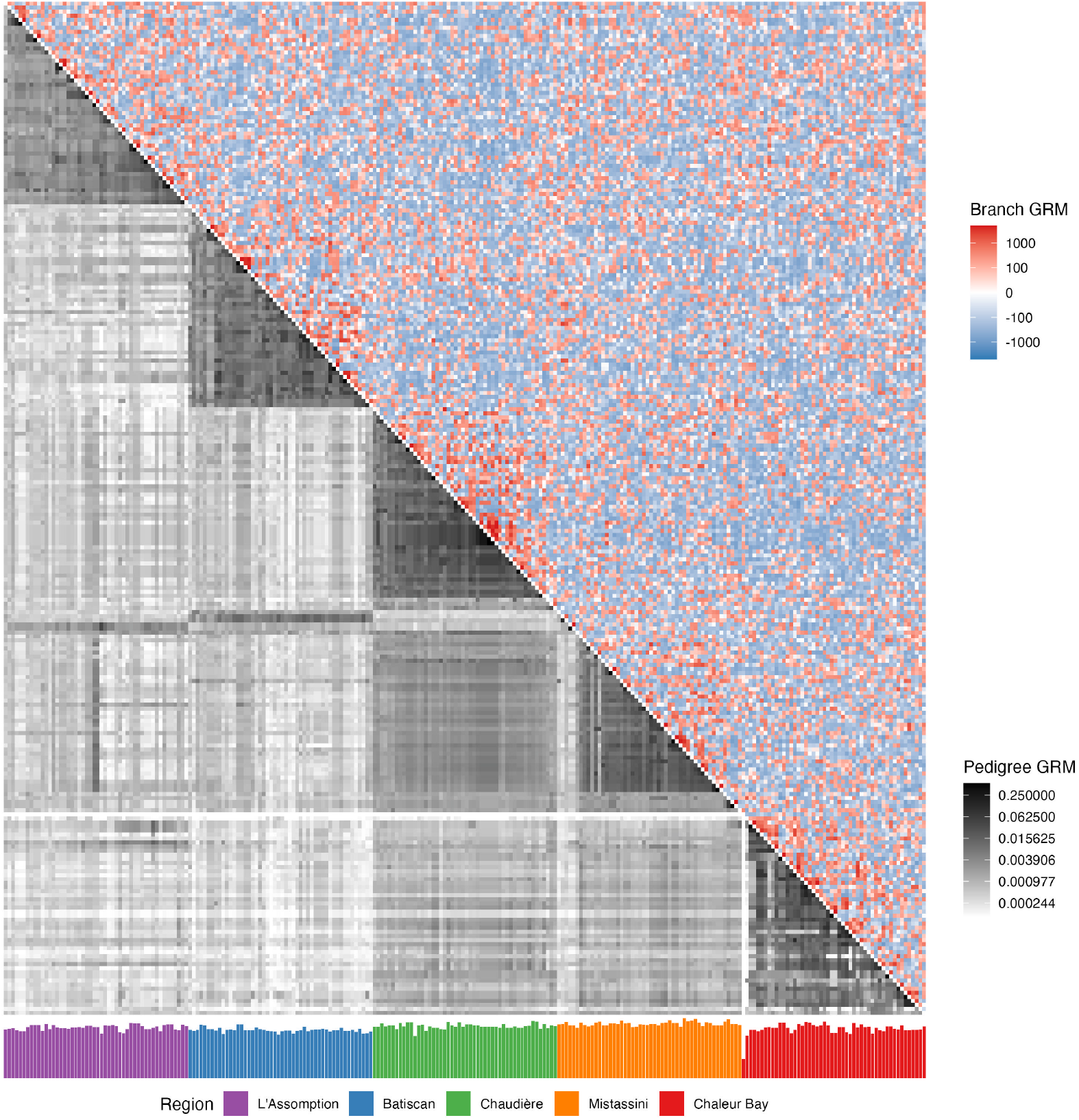
Relatedness between 250 French-Canadian individuals from 5 regions in Quebec. Upper-triangle: Heatmap of the branch GRM computed from one ARG (one chromosome). Lower-triangle: Heatmap of the pedigree GRM. Bottom: Barplot of the average founder depth in the pedigree for each individual. The ordering of individuals is based on region and within region hierarchically on pedigree GRM. Because of the log scaling in the heatmap of branch GRM, we used an epsilon of 10^−4^ to avoid issues with values close to zero.

Next, we explored how the branch relatedness of a given pair of individuals varies across 100 ARGs simulated within the pedigree, for a large number of pairs (Figure 3). As expected, branch relatedness increases with pedigree relatedness. Since pedigrees are often shorter than the mean TMRCA, and so the contribution of branch lengths within the pedigree is small, a simple approximation of the relationship between the two can be derived as follows. First, since pedigree relatedness *r*_*i,j*_ between a pair of individuals is the expected proportion of the genome on which the two inherit from a common ancestral genome within the pedigree, we expect branch relatedness *C*_*i,j*_ to be roughly *r*_*i,j*_*C*_0_ + (1 − *r*_*i,j*_)*C*_∗_, where *C*_0_ is the average branch relatedness of a genome to itself in pedigree founders, and *C*_∗_ is the average branch relatedness of two distinct genomes from the pedigree founders. Since this relatedness is *centered*, we can (very roughly) take *C*_∗_ ≈ 0 and *C*_0_ ≈ *A*(*U, U*) − *A*(*U, V*) (with *U* and *V* random individuals). In other words, the average centered branch relatedness of a typical genome to itself is the total area of edges back to the roots, minus shared edges between two typical (but different) genomes. Using the relationship (13), this is *C*_0_ ≈ *D*(*U, V*) /2. Hence, we expect **B**_*i,j*_ ≈ *r*_*i,j*_*T* where *T* is the mean TMRCA for two random samples from the population (computable as one half of branch genetic divergence in `tskit`). This is shown with a line in Figure 3, with *T* computed from the demographic model used for recapitation of the pedigree.

**Figure 3:**
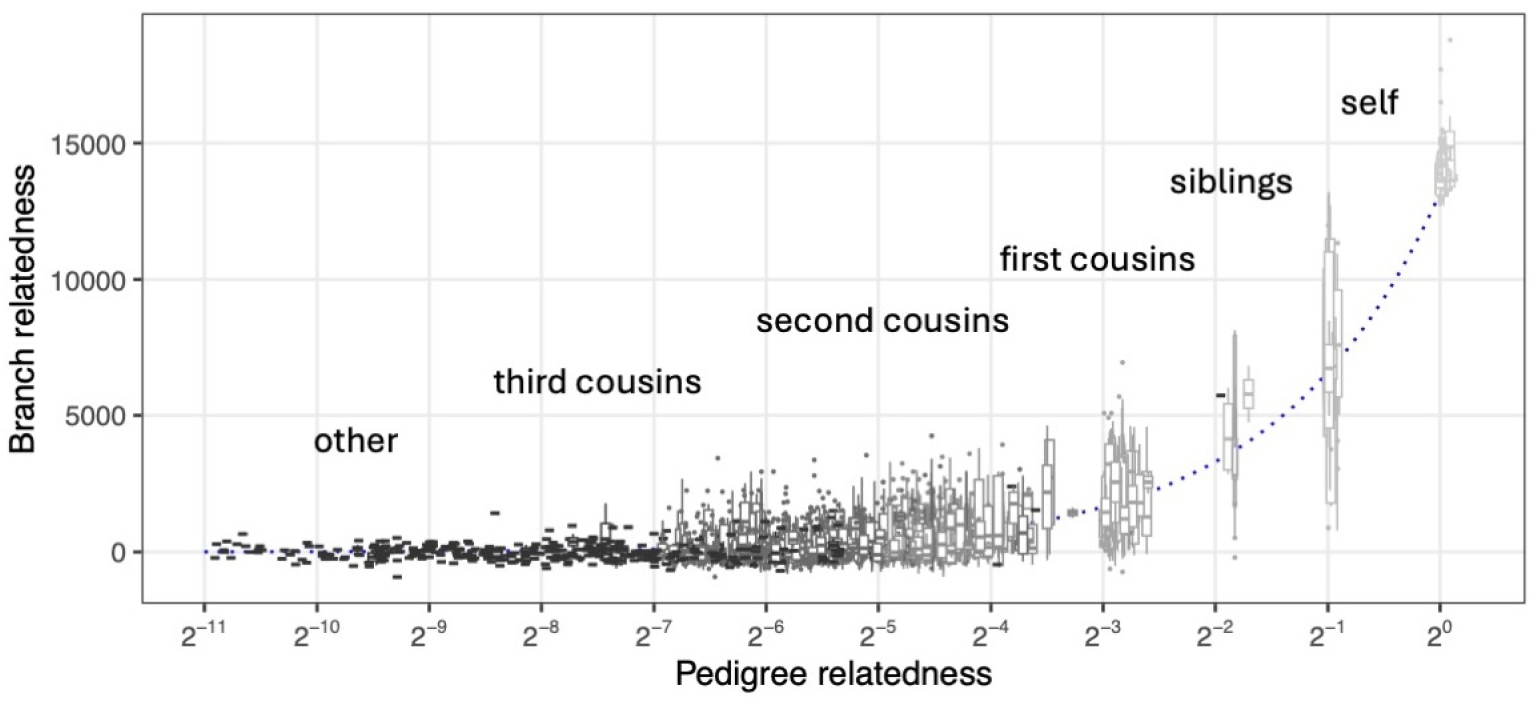
Variability in branch relatedness with respect to a fixed pedigree. Each box plot corresponds to a pair of individuals with pedigree relatedness according to some types of (pedigree) relationships (self, siblings, etc.). The box plot for each pair of individuals depicts variation in branch relatedness across 100 ARGs within the fixed pedigree. The dotted line indicates the approximate expected branch relatedness, which is the pedigree relatedness multiplied by the mean TMRCA among pedigree founders.

However, there is a degree of variability in branch relatedness between different pairs of individuals with similar pedigree relatedness. While branch relatedness broadly tracks the expected relatedness outlined above, it varies around this value across the range of pedigree relatedness. Moreover, there is substantial variability in branch relatedness across one chromosome for a pair of individuals. This variability is highest for sibling pairs and decreases with pedigree relatedness. We expect the absolute variability to decrease when considering branch relatedness across the whole genome.

### 2.7 Computation

We next present efficient algorithms for various operations, which are implemented in the `tskit` library [Ralph et al., 2020, Kelleher et al., 2024].

#### 2.7.1 Computation of the entire branch GRM

As shown in equation (14), the branch GRM **B** is a straightforward function of the more fundamental divergence matrix **D**. The divergence matrix describes the total branch length separating all 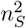 pairs of samples, such that **D**_*i,j*_ = 2*t*_*u*_ − *t*_*i*_ − *t*_*j*_ where *u* is the MRCA of samples *i* and *j*. Because the output is a dense *n*_*S*_ × *n*_*S*_ matrix and at a minimum we must create and fill in the entries of this matrix, the complexity of this operation is at least 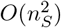.

So-called “incremental algorithms”, which use the fact that the small changes in tree structure we observe due to recombination events often correspond to small changes in some accumulated statisic as we move along the genome, have led to very efficient algorithms in several contexts [Kelleher et al., 2016, Ralph et al., 2020, Kelleher and Lohse, 2020]. The divergence matrix, however, does not easily lend itself to this approach. Incremental algorithms work well when we only need to consider the effects of inserting and removing edges on nodes that are *ancestral to* a given node. To compute the divergence matrix, however, we need to keep track of when the MRCAs of each pair of samples change, and this requires traversing the subtrees *descending from* nodes affected by edges being inserted and removed. Removing (or inserting) an edge changes the MRCA of all pairs between the set of samples descending from it and those not descending from it. In worst case (removing an edge to the root of a balanced binary tree with *n*_*S*_ samples) this involves 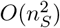 work per tree transition, and therefore the complexity of the operation is 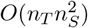, where *n*_*T*_ is the number of trees along the genome.

The naïve approach to this problem is to proceed tree-by-tree along the sequence, iterate over all 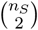 pairs of samples, compute the time to their MRCA, and update the corresponding element of **D**. MRCAs can be computed efficiently using the Schieber-Vishkin algorithm [Schieber and Vishkin, 1988, *Knuth, 2011] which provides the MRCA of two nodes in constant time after an O*(*n*_*S*_) preprocessing step. The overall complexity is therefore 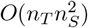, as we need to perform 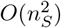 work for all *n*_*T*_ trees. While this is the same complexity as the incremental approach outlined above, this “naïve” approach is in practise much faster, and is therefore the implementation used in `tskit` via the `ts.genetic_relatedness_matrix` method. The `eGRM` package Fan et al. [2022] essentially uses the same approach, although implemented in Python and without efficient bulk MRCA queries. Their approach therefore requires 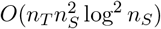 time, as each MRCA query requires *O*(tree height) time, which is log *n*_*S*_ if the trees are balanced [Kelleher et al., 2016]. Figure A1 (Appendix D) shows that the `tskit` implementation is faster than the `eGRM` implementation [Fan et al., 2022], although converging for larger sample sizes.

The 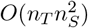 complexity of computing the branch GRM has significant implications for its utility in large-scale studies: quadratic algorithms are simply not feasible when we have millions of samples. The approximate mutation-dropping approach of Zhang et al. [2023] is not directly comparable to Fan et al. [2022] and our work. However, their follow-up work with the randomized Haseman-Elston method [Zhu et al., 2024] indicates that there are scaleable computational approaches that can work with approximate branch GRMs. In the next section, we show that it is not necessary to compute the exact branch GRM explictly in order to *use* it. For many applications that use the branch GRM we can implicitly compute with it without materialising the actual matrix.

#### 2.7.2 Computing branch GRM-vector products

Many calculations with GRMs involve matrix-vector products [Colleau, 2002, Colleau et al., 2017]. The straightforward way is to first compute the GRM, and to then compute its product with the vector in question. This places strong limits on the size of the datasets that we can work with, due to quadratic space and time complexity in the number of samples, *n*_*S*_. In certain situations, however, given that the output of an (*n* × *n*)-matrix times an *n*-vector is, itself an *n*-vector, we can perform this calculation without explicitly materialising the matrix, thus avoiding the quadratic space complexity. In rarer situations, we can even exploit the structure of the matrix to also avoid the quadratic *time* complexity. Not only do we not fully materialize the matrix in memory, we never actually compute all 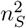 elements of the matrix. Here, we develop an algorithm of the latter form, which allows us to exactly compute the product of the branch GRM with an arbitrary vector in substantially less time than it would take to compute the matrix itself. Roughly, this works because although the GRM is not itself sparse or low rank, the ARG provides a decomposition of the ARG into a sum of low-rank components with hierarchical structure – the (sub)trees.

To do this we require some notation. Suppose now that **C**_*s,t*_ is the uncentered branch relatedness between sampled genomes *s* and *t* as computed from the trees, that is, the sum of the areas of all branches in all trees that are ancestral to both *s* and *t*. This is **C**_*s,t*_ = ℂov[*Z*(*s*), *Z*(*t*)] as in equation (7). For a given vector **w**, we’d like to compute the matrix-vector product **Cw**. Write *b*_*k*_ (*k* = 0, …, *K*) for the unique recombination breakpoints on the genome including the start and end of the genome (the genome is a closed interval r*b*_0_, *b*_*K*_s). Suppose that the *k*^th^ tree *T*_*k*_ extends over the region from *b*_*k*_ to *b*_*k*−1_ along the genome, that the length of the edge (in units of time) above node *n* in *T*_*k*_ is 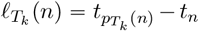, where 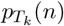 is the parent node of *n* in *T*_*k*_. Finally, write ≤_*T*_ for the partial ordering of nodes induced by tree *T* with older nodes larger than younger nodes. Then, the uncentered branch relatedness matrix is

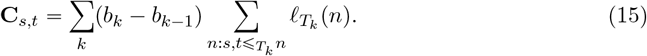

The *s*^th^ element of the matrix-vector product **Cw** is:

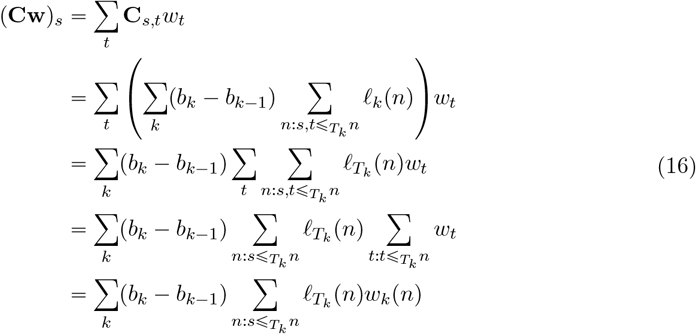

Here 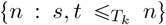 is the set of nodes *n* that are ancestral to both *s* and *t* in *T*_*k*_, and 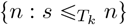 are those nodes just ancestral to *s*. The new variable *w*_*k*_(*n*) is the sum of sample weights below *n* in tree *T*_*k*_:

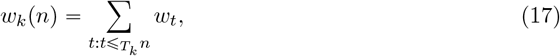

which is a familiar term from Ralph et al. [2020]. Although a single entry of **Cw** could be computed efficiently from the algorithm in Ralph et al. [2020], it doesn’t scale well because it requires a separate set of weights for each entry of the vector.

We present an efficient algorithm for computing the entire matrix-vector product. The general idea is simple: as we move left-to-right along the tree sequence, we keep track of two things for each node *n*: the *weight w*(*n*) of the node in the current tree (*w*_*k*_(*n*) above) and the *value v*(*n*) of the haplotype carried by *n*, which will contribute to all descendants of *n*. Additionally, we keep track of the last *position x*(*n*) in which the node was updated. As we move along the genome, we update any nodes ancestral to any changes in the tree: all other nodes are the roots of unchanged subtrees and hence remains unchanged. As seen above, each branch contributes to potentially many entries in the output vector, so by accumulating values of haplotypes, we reduce the amount of necessary work.

##### Algorithm V

(*Branch GRM-vector product*). Given a sequence of positions that are recombination breakpoints *b*_*k*_ for 1 ≤ *k* ≤ *K* along the genome and corresponding sequences of edges to remove (*R*_*k*_) and add (*A*_*k*_) at each position, compute the values *y*_*s*_ = ∑_*t*_ **C**_*st*_*w*_*t*_ for 1 ≤ *s* ≤ *n*_*S*_, assuming all samples are leaves in all trees. Let *T* be the current tree, 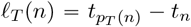 be the length of the branch above *n* in *T* (or zero, if *n* has no parent), initialize *k* = 1, *x*(*n*) = 0, and *v*(*n*) = 0 for all *n* ∈ **V**. Set *w*(*s*) = *w*_*s*_ for each sample *s*, and *w*(*n*) = 0 for all other nodes. Let *z*(*n*) = ℒ_*T*_ (*n*) (*b*_*k*_ − *x*(*n*)) be a function computed from the current values of *k* and *x* at all times.

**V1**. [Remove edges] For each edge (*c, p*) ∈ *R*_*k*_, and for each node *n* ≥_*T*_ *p*, set *v*(*n*) + = *z*(*n*)*w*(*n*), then *w*(*n*) − = *w*(*c*), *v*(*c*) + = *v*(*n*), and *x*(*n*) = *b*_*k*_. Then, set *x*(*c*) = *b*_*k*_ and remove the edge.

**V2**. [Add edges] For each edge (*c, p*) ∈ *A*_*k*_, and for each node *n* ≥_*T*_ *p*, set *v*(*n*) + = *z*(*n*)*w*(*n*), then *w*(*n*) + = *w*(*c*), *v*(*c*) − = *v*(*n*), and *x*(*n*) = *b*_*k*_. Then, set *x*(*c*) = *b*_*k*_ and add the edge.

**V3**. [Iteration] If *k* < *N*, set *k* + = 1 and return to **V1**. Otherwise, set *y*_*s*_ = *v*(*s*) for 1≤ *s* ≤ *n*_*S*_ and finish.

Algorithm V follows a similar structure to previous incremental algorithms [Kelleher et al., 2016, Ralph et al., 2020]: at each tree transition we update some global state to account for the insertion and removal of the edges affected. Here, the overall goal is different: rather than keeping track of some cumulative value among the nodes in a given subtree (say, total branch length) we are instead keeping track of the total contribution to each node from nodes *ancestral to it*. By some subtle bookkeeping, we can keep track of the cumulative contribution to each node, in only updating each node when it is affected by an edge insertion or removal. Each node accumulates the contributions that are passed down from above until an edge below it is added or removed. At each edge insertion or removal *v*(*n*) is updated by traversing up to the root of the current subtree (also keeping the weights *w*(*n*) up to date), and the accumulated contribution passed down to the child node of the edge *c*. Finally, we set *x*(*c*) is set to *b*_*k*_ (the current position) to mark the last position this node was updated.

The above explanation is a rough sketch of the algorithm. A full proof of correctness is provided in Appendix E. The algorithm has been implemented in the `ts.genetic relatedness vector` method in `tskit`, somewhat generalized to allow for samples that are not leaves, and is extensively tested.

The analysis of this algorithm is straightforward and follows a standard pattern [Kelleher et al., 2016, Ralph et al., 2020]. Because recombination results in a small modification of the current tree, each tree transition incurs *O*(1) edge removals and insertions. Each edge removal in step **V1** involves examining only nodes ancestral to the edge, and therefore incurs a cost of *O*(log *n*_*S*_), assuming trees are balanced. Edge insertions in **V2** have the same cost. Thus, as the first tree requires inserting *O*(*n*_*S*_) edges requiring *O*(1) work, the overall complexity is *O*(*n*_*S*_ + *n*_*T*_ log *n*_*S*_). This logarithmic time complexity is borne out in Figure 4 where we plot the time taken to compute the branch GRM-vector product against subsets of a large simulated ARG [Anderson-Trocmé et al., 2023]. Here, it takes only 17.8 seconds to run the `ts.genetic relatedness vector` method on the ARG with 1 million diploid samples (6,694,080 nodes; 31,840,754 edges; 4,013,273 trees). In contrast, computing the full branch GRM using the `ts.genetic_relatedness_matrix` method for the ARG with ten diploid samples (61,412 nodes; 297,171 edges; 93,543 trees) required 28 seconds.

**Figure 4:**
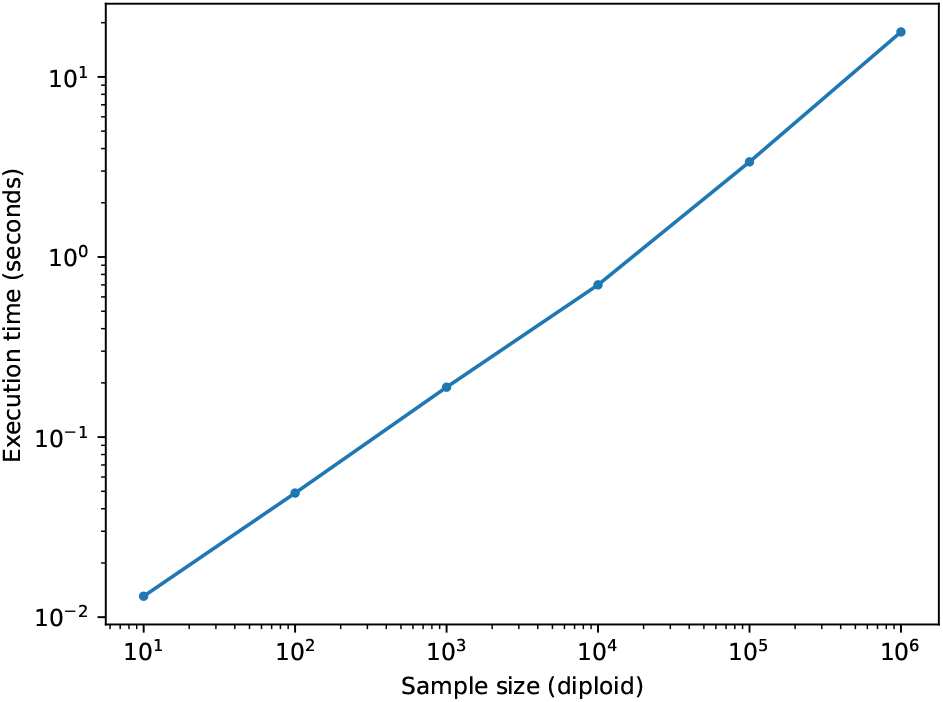
Computational scaling of the branch GRM-vector product algorithm implemented in tskit for subsets of a large simulation of French-Canadians.

#### 2.7.3 Branch PCA

We found the principal components (PCs) of the branch GRM using a randomized SVD [Halko et al., 2011], a method that can find the eigenvectors of a matrix that is only implicitly defined through a matrix-vector multiplication. We implemented the algorithm as `ts.pca` in `tskit`.

##### Algorithm rPCA

(*Randomized PCA of branch GRM*). Let **C** be the branch GRM for *n*_*I*_ individuals, let *k* be the desired number of PCs, and *q* the number of iterations. Multiplying **C** with a vector is done by Algorithm **V**.

**P1**.[Range estimation] Sample a random matrix 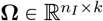 in which the entries are independent standard normal variables. Obtain a basis matrix 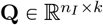 by applying QR decomposition to **CΩ**. Repeat *q* times, updating the basis matrix **Q** by applying QR decomposition to **CQ**, where **Q** is from the previous iteration.

**P2**.[Small singular value decomposition] Compute **W** = **Q**^**⊤**^**C** and obtain the singular vector **U** ∈ R^*k*×*k*^ by exact singular value decomposition of **W**. Then the columns of **QU** P R^*n*×*k*^ contain the desired PCs of the branch GRM **C**.

The algorithm has two advantages over directly applying the exact SVD to the branch GRM. It needs less time and memory because the *n*_*I*_ × *n*_*I*_ branch GRM is never computed nor stored. The algorithm extracts the relevant information through the efficient matrix-vector product Algorithm V. Secondly, the exact SVD is applied to an *n*_*I*_ × *k* matrix, where *k* is much smaller than *n*_*I*_. This reduces the amount of computation considerably.

The efficiency of the branch PCA algorithm and the underlying branch GRM-vector product algorithm is illustrated in Figure 5. See Methods for details of the benchmarking methodology. For PCA, we observed significant benefits from using implementations that avoided the storage of the GRM or genotype matrix in memory, particularly for larger numbers of samples (Figure 5). Notably, `ts.genetic_relatedness_matrix` failed due to memory limits when computing the branch GRM for 2^12^ = 4096 sample nodes and when computing the genotype (site) GRM for 2^14^ = 16384 sample nodes. Randomized PCA on the genotype matrix in `scikit-allel` failed due to memory limits for 2^16^ = 65536 sample nodes. Implementations that relied solely on the implicit matrix-vector product using `tskit` were substantially more efficient: both `ts.pca` and `eigsh` from `scipy` using `ts.genetic relatedness vector` as a linear operator were able to scale to 2^20^ = 1, 048, 576 samples. The native implementation of `ts.pca` consistently outperformed `eigsh`, with the relative difference decreasing slightly with the number of samples, and increasing with sequence length. Moreover, the difference in absolute compute time increased with both sample size and sequence length. For example, `ts.pca` took on average 0.27s for 2^12^ = 4, 096 samples and 26.9s for 2^20^ = 1, 048, 576 samples, while `eigsh` took 1.7s for 2^12^ = 4, 096 samples and 119.7s for 2^20^ = 1, 048, 576 samples. This difference primarily reflects the differences in the underlying algorithms used for PCA: `ts.pca` uses a randomized SVD while `eigsh` uses the implicitly restarted Lanczos method [Lehoucq et al., 1998].

**Figure 5:**
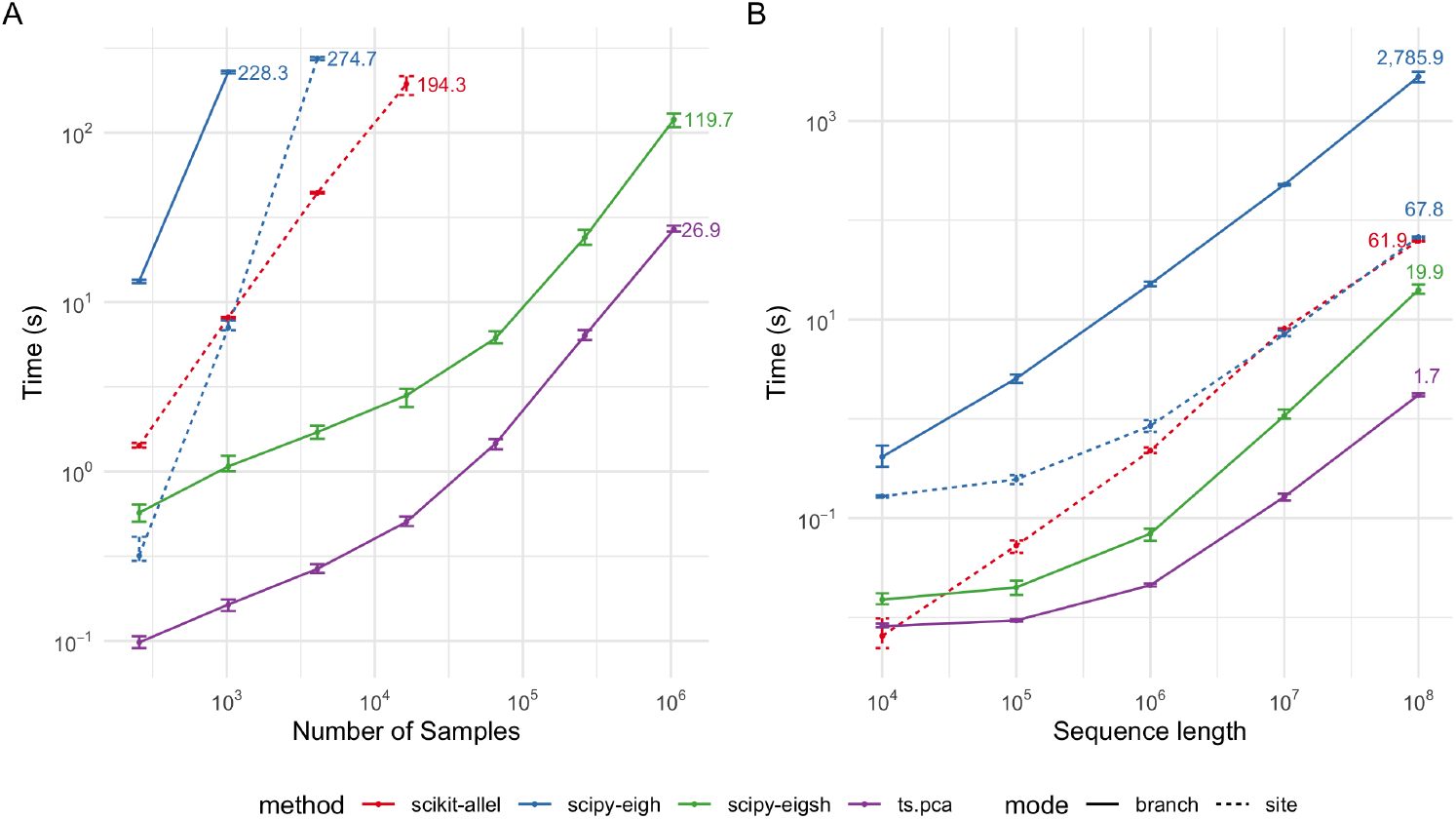
Time efficiency of different implementations of PCA computations. Each dot corresponds to the average time taken across ten simulations with different random seeds. Error bars represent the range in time taken across the ten simulations. (A) PCA with genome sequence length fixed at 10^7^ and varying the number of samples. (B) PCA with number of sample nodes fixed at 2^10^ and varying genome sequence length. Branch mode refers to branch PCA and site mode refers to genotype PCA.

## 3 Methods

### 3.1 French-Canadian pedigree

To demonstrate the similarities and differences between pedigree and branch relatedness, we performed a range of analyses on a subset of an extended pedigree of French-Canadian individuals from the BALSAC project [Vézina and Bournival, 2020]. Spanning over 400 years, this pedigree is compiled from over 4.5 million parish records across Quebec. In this paper, we restricted our analyses to five regions, each containing five neighboring parishes (Table 1). Using the migratory patterns from Anderson-Trocmé et al. [2023] as a reference, we identified five distinct regions and sampled individuals from parishes within each region to minimize excessive relatedness within each sampling unit while also mitigating the risk of de-anonymization. Our data access agreements with BALSAC dictated that we use parish records of more than one hundred years old (before 1924) to publish their metadata and summary statistics. As a result, the pedigree used in this study contains ascending genealogy for 500 randomly chosen contemporary individuals from each of the five regions, with individuals sampled across the five selected parishes per region.

**Table 1:** Selected regions and parishes from the BALSAC French-Canadian Pedigree

| Region | Parishes | Approximate location |
| --- | --- | --- |
| Chaleur Bay | St Michel, St François De Sales, St Georges De Malbaie, St Pierre De Malbaie, and St Joseph | 48.5222, -64.2156 |
| Batiscan | Ste Geneviève De Batiscan, St Luc De Vincennes, St Narcisse, and St Stanislas | 46.5324, -72.3398 |
| Chaudière | St Georges, St Benoit Labre, St Philibert, St Come, and St Martin De Tours | 46.1184, -70.6691 |
| L’Assomption | St Jacques, St Alexis, Ste Marie Salomé, and St Esprit | 45.9483, -73.5702 |
| Mistassini | St Félicien, St Méthode, Notre Dame De La Dore, St Cyrille, and Ste Lucie | 48.6399, -72.4543 |

The sub-pedigree obtained from the selected five regions, each containing five parishes, consisted of 61,490 individuals, including 2,321 probands. A subset of the probands exhibited a low depth of pedigree, reflecting an incomplete pedigree. To ensure meaningful comparisons, we computed the maximum pedigree depth for each individual and derived an average depth metric. A total of 48 probands with an average depth of less than 3 were excluded due to their shallow pedigree. After this filtering step, a total of 2,273 probands were retained for downstream analyses.

We computed pedigree and branch relatedness between individuals of interest. Pedigree relatedness was computed after Lange and Sinsheimer [1992] and Colleau [2002]. Branch relatedness was calculated from an ARG obtained with the simulation based on pedigree and ancestry described in Anderson-Trocmé et al. [2023]. In short, this simulation uses `msprime` [Baumdicker et al., 2022] for a backward in time simulation in two stages. First, it samples chromosomal inheritance through the fixed pedigree to obtain an ARG within the pedigree. Second, it simulates the ancestry of the ARG obtained in the first stage by coalescent simulation from a given demographic model (i.e., “recapitation”). We elected to simulate only the complete human chromosome 3 due to its large size while reducing the overall computational cost of our study. To study the stochastic variation in recombination and coalescence events within the pedigree, we simulated 100 replicates of an ARG for the chromosome using different random seeds. This approach allowed us to assess the variance in branch GRM while maintaining consistency with the underlying pedigree.

To explore the overall population structure within the pedigree and the simulated tree sequences, we performed PCA on the set of 2,273 probands. For pedigree PCA, we first computed the pedigree GRM among the probands and then eigen-decomposed the GRM using `eigh` function from `scipy` [Virtanen et al., 2020]. For branch PCA, to avoid undue influence of large, low-recombination regions, we first remapped genomic coordinates from base pairs to genetic distance, using the HapMap II genetic map provided by `stdpopsim` [Adrion et al., 2020]. We then used Algorithm rPCA to compute the first six PCs.

To compare pedigree and branch relatedness for specific pairwise relationships, we focused our attention on a subset of the 2,273 probands. Specifically, we randomly sampled one parish per region and subsampled at least five siblings, first cousins, second cousins, and third cousins from each parish. We then subsampled additional individuals from each parish to obtain a total of 50 individuals per parish. With these 250 individuals, we computed the pedigree GRM and a branch GRM for each of the 100 simulated ARGs.

### 3.2 Benchmark simulations

We assess the computational efficiency of our implementations for branch GRM and PCA, with simulations, recording the time for calculations for a range of tree sequences. We simulated the tree sequences with `msprime` [Baumdicker et al., 2022], and varied either the genome sequence or the number of sample nodes (=haploid individuals). All computations were carried out on a single CPU with 4GB of RAM.

#### 3.2.1 Branch GRM

We compared `ts.genetic_relatedness_matrix` for computing the branch GRM to the implementation in the `eGRM` package [Fan et al., 2022]. The default values for simulations parameters were 10^7^ for the genome sequence length, 2^10^ for the number of samples, and effective population size of 10^4^. We then varied genome sequence length from (10^4^, 10^5^, 10^6^, 10^7^, 10^8^) and the number of samples from (2^7^, 2^8^, 2^9^, 2^10^, 2^11^, 2^12^), each one at a time. For each simulation setting, we generated 10 tree sequences with different random seeds and reported the average time taken to compute the GRM with each implementation.

#### 3.2.2 Branch PCA

We assessed our branch PCA Algorithm rPCA against a number of comparators using `scipy` [Virtanen et al., 2020] and `scikit-allel` [Miles et al., 2024]:

- Calculating branch GRM using `ts.genetic_relatedness_matrix` followed by eigenanalysis using `eigh` function from `scipy`.
- Eigenanalysis of branch GRM using `eigsh` function from `scipy` using the implicit matrixvector product Algorithm V as a linear operator.
- Calculating genotype GRM using equation (4) followed by eigenanalysis using `eigh` function from `scipy`.
- Randomized PCA of genotype matrix using `randomized pca` function from `scikit-allel`.

We used the same simulations as for the branch GRM computation benchmark, but explored larger sample sizes, ranging across (2^8^, 2^10^, 2^12^, 2^14^, 2^16^, 2^18^, 2^20^). For each simulation setting, we generated 10 tree sequences with different random seeds and reported the average time for PCA with each implementation.

## 4 Discussion

Recent advances in ARG inference have generated significant interest in leveraging ARGs for genetic analyses. In this paper, we examined the relationship between different definitions of genetic relatedness in the common context of additive traits on an ARG, especially the emergent notion of “branch” relatedness. We also demonstrated how branch relatedness compares with pedigree relatedness in simulations through a pedigree of French-Canadians. We then described an algorithm for branch GRM-vector product, to bypass the fundamental problem of quadratic complexity of computing and storing GRMs. This algorithm allowed us to use randomized linear algebra methods for branch PCA using an ARG on a million samples in 30 seconds and less than 4GB of RAM. We close the discussion with an outlook for the use of these algorithms in population and quantitative genomic analyses.

The described branch relatedness unifies several notions of relatedness into one framework by leveraging the ARG encoding of how sampled genomes relate to each other. One thing that distinguishes these notions of relatedness is which aspects of genetics and genealogy are unobserved or observed, and which are averaged over or fixed. For instance, pedigree relatedness averages over recombinations within a known pedigree, genotype relatedness averages over typed loci stored in a genotype matrix, while branch relatedness averages over mutations on inherited genome segments in an ARG. Our branch relatedness is conceptually equivalent to the eGRM from Fan et al. [2022] and the ARG-GRM from Zhang et al. [2023], although we omit the scaling by (*p*(1 − *p*))^*α*^ used by both. The “e” in eGRM denotes expectation of the genotype GRM under a Poisson mutation process along ARG branches. At the risk of further confusing terminology, we adopted the term *branch* to highlight that this measure of similarity is derived from the extent of shared branch area between individuals, explicitly distinguishing it from expectations that are conditional on other quantities – for instance, expected covariance given a pedigree or expected covariance given a collection of genotypes (but not their effect sizes). We have seen that branch relatedness varies substantially for relatives of a given degree, in line with the theory on pedigree and genetic ancestors [Chang, 1999, Weir et al., 2006, Hill and Weir, 2011, Thompson, 2013, García-Cortés et al., 2013]. Does mutation contribute a large degree of variability in addition to the branch relatedness? Fan et al. [2022] computed the “varGRM” to describe this (for general theory see Ralph [2019]); and in general the answer is “no” – randomness due to the mutation process adds little variability beyond recombination, except to small segments of the genome.

We have chosen to interpret GRM in the context of a generative model of traits following the initial definition of relatedness [Wright, 1922]. However, the worth of a given GRM is determined by how well it works in practice, rather than its theoretical justification, and applications have motivated a number of interpretations and adjustments [Speed and Balding, 2015]. However, adjusting the trait model gives a natural setting in which to suggest extensions and the corresponding GRM is then a by-product of these extensions. Although our current definition of relatedness assumes equal prior effects across all loci, one could consider alternatives whereby we incorporate prior information on effect sizes. For example, selection reduces frequency of deleterious mutations with strong effects from the population and such mutations may justify a different prior; this prior might depend on mutation age in a similar way that the GRMs often weigh alleles by a function of their frequency [Speed and Balding, 2015]. Functional annotations have been used to improve fine-mapping and genomic prediction [e.g. MacLeod et al., 2016, Weissbrod et al., 2020, 2022] and could be incorporated as prior information on mutation effects, which will refine branch relatedness calculations for trait-based analyses.

Computing a full GRM is inherently a quadratic operation and therefore not feasible on large sample sizes. It is possible, however, to calculate GRM-vector products at a substantially lower computational cost. With *n*_*S*_ samples and *n*_*T*_ local trees, our branch GRM-vector product algorithm has complexity *O*(*n*_*S*_ + *n*_*T*_ log *n*_*S*_). This relies on two insights: first, we use local trees to efficiently encode the low-dimensional block structure of the contribution to the GRM of a single local tree; and second, we leverage the fact that most tree structure is shared across many local trees in the ARG. This removes the need for approximate methods such as the Monte Carlo sampling of mutations on the ARG used by Zhang et al. [2023]. The method is most similar to Zhu et al. [2024], who uses iterative algorithms to compute GRM-vector multiplication with the genotype GRM from Monte Carlo-sampled mutations, but the algorithm is not given, so a more detailed comparison is not possible. We also provide highly efficient vector-GRM-vector product algorithm, similar to the classic algorithm for large pedigrees [Colleau, 2002], using the generic framework of Ralph et al. [2020].

This work provides the definition of branch relatedness based on a concrete trait model, algorithms to efficiently compute with the corresponding branch GRM for millions of genomes, and well documented and thoroughly tested open-source `tskit` implementation. These contributions are further opening a possibility for mega-scale population genetics and quantitative genetics. The clear definition of branch relatedness (based on the fundamental ARG encoding of sampled genomes, a trait model extendable with additional biological prior information) will enhance the analyses of diverse and admixed genomic datasets that are challenged by many evolutionary processes and data availability [e.g. MacLeod et al., 2014, Martin et al., 2017, Durvasula and Lohmueller, 2021, Wang et al., 2022, Yair and Coop, 2022, Ros-Freixedes et al., 2022a]. The efficient branch GRM-vector product algorithm will speed-up analyses of population structure, genome-wide associations, heritability, and genomic prediction.

## Data Availability

We thank the BALSAC project for providing access to their genealogical data and for their guidance in selecting an appropriate subset of the genealogy for our analyses. Contact BALSAC for more information and to apply for access to these data (https://balsac.uqac.ca/)

## Acknowledgments

We are grateful to Georgia Tsambos, Yan Wong, Nate Pope and John Novembre for helpful discussions and comments on the manuscript. The authors acknowledge the use of the UCL Myriad High Performance Computing Facility (Myriad@UCL), and associated support services, in the completion of this work. Luke Anderson-Trocmé was supported by R35 GM149521 and NSERC PDF-588001-2024. Gregor Gorjanc acknowledges support from the BBSRC ISP grant to The Roslin Institute (BBS/E/D/30002275, BBS/E/RL/230001A, and BBS/E/RL/230001C) and BBSRC research grant (BB/T014067/1). Jerome Kelleher and Peter Ralph were supported by R01 HG012473 from the National Institutes of Health NHGRI. Jerome Kelleher acknowledges support from EPSRC (research grant EP/X024881/1) and the Robertson Foundation. Hanbin Lee acknowledges support from the Statistics Department at the University of Michigan through the Departmental Fellowship. Brieuc Lehmann acknowledges support from the UKRI [EP/R018561/1; UK Engineering and Physical Sciences Research Council (EPSRC) Bayes4Health programme] and funding from Jesus College, Oxford.

### Box 1: A Brief History of Genetic Relatedness

Genetic relatedness was first formalised by Wright [1922], who introduced the coefficients of relationship and inbreeding in the context of a pedigree based on the path (correlation) analysis of phenotypic values for pedigree members. He also showed how to compute these coefficients in a general pedigree by tracing all pedigree lineages between relatives. Emik and Terrill [1949] and Cruden [1949] devised a simpler procedure to compute these coefficients between all pairs of pedigree members, which was later formalised as a recursive algorithm [Henderson, 1976]. The algorithm fills in a symmetric genetic relatedness matrix (GRM), which is the key object for statistical genetics, particularly through its use in linear mixed models [Falconer and Mackay, 1996, Henderson, 1984, Lynch and Walsh, 1998, Mrode and Pocrnic, 2023]. However, pedigree information is limited because it can only quantify the relatedness relative to pedigree founders [Wright, 1965, Jacquard, 1975], pedigrees are increasingly incomplete backwards in time [e.g. Legarra et al., 2015], and pedigrees measure expected rather than realized relatedness due to recombination and mutation [e.g. Hill and Weir, 2011, Thompson, 2013, García-Cortés et al., 2013].

Relatedness was later redefined with respect to genotypes using concepts of identity by descent (IBD) and identity by state (IBS) by Cotterman [1940] and Malécot [1948, 1969], before DNA data was readily available. Early developments of molecular genetics enabled generation of DNA data and estimation of genotype-based relatedness (see review by Weir et al. [2006]). As anticipated by Thompson [1975], early studies found that the number of markers is critical for the accuracy of estimates and distinguishing different types of relationships for pairs of individuals. Increasing the number of markers improved the accuracy, but also revealed substantial variation from the expected pedigree relatedness between pairs of individuals - overall as well as along genome regions - due to recombination [Weir et al., 2006].

Further developments in molecular genetics streamlined genome-wide genotyping with SNP arrays and increasingly whole-genome resequencing such that today we have genotype datasets with thousands to millions of individuals [e.g. Turnbull et al., 2018b, Bycroft et al., 2018, Ros-Freixedes et al., 2022b]. This data-abundance has reinvigorated population genetics studies of variation within and between populations [Begun et al., 2007, Langley et al., 2012], and statistical genetics studies of complex trait architecture [Burton et al., 2007, Abdellaoui et al., 2023] and prediction of such traits [Meuwissen et al., 2001, 2013]. Depending on the aims of the study [Speed and Balding, 2015], we now use a number of different relatedness estimators [e.g. VanRaden, 2008, Yang et al., 2010, Manichaikul et al., 2010, Speed et al., 2012, Weir and Goudet, 2017, 2018, Ochoa and Storey, 2021, Mary-Huard and Balding, 2023]. Variants of the genotype sample covariance between individuals, known as the genomic relationship/relatedness matrix (GRM), are commonly used [VanRaden, 2008, Yang et al., 2010, Speed et al., 2012]. These estimators largely treat loci independently, with research on leveraging linkage between loci to improve delineation between IBS and IBD relatedness [e.g. Visscher et al., 2006, Browning and Browning, 2012, Thompson, 2013, Hickey et al., 2013, Edwards, 2015, Pook et al., 2019, Nait Saada et al., 2020]. The ease of use is the primary reason for treating genotype loci independently. Conversely, to leverage linkage between loci, one needs to phase genotypes, operationally define similarity between the resulting haplotypes, and then estimate relatedness based on these haplotype similarities.

There is a growing interest to use ancestral recombination graphs (ARG) as the ultimate description of such haplotype similarities for a sample of individuals. ARGs are therefore enabling the study of relatedness and its use in downstream genetic analyses [Fan et al., 2022, Tsambos, 2022, Zhang et al., 2023, Link et al., 2023, Schraiber et al., 2024]. Relatedness based on an ARG leverages allele differences between individuals, but also linkage between these alleles, hence connecting IBS and IBD concepts, capturing typed and untyped loci, and time-varying changes in population structure due to ancient and recent demographic events [Fan et al., 2022, Young, 2022, Zhang et al., 2023, Harris, 2023].

## Appendix

### A Treatment of reference alleles

In the model above we have set the effect of a given allele (either the reference, major, or ancestral allele) to zero, and assigned the effect of each other allele an independent Gaussian effect. This, however, apparently depends on the choice of reference allele, and so another choice would have been to assign a separate independent Gaussian effect to *every* allele. It turns out that this is only equivalent in the biallelic case, in the sense that there exists a transformation of parameters for one model that produces the other model. The situation is essentially that of choosing the contrasts for a factor when fitting a linear model with Gaussian priors on the parameters: the BLUPs are the same, but the posterior distributions on the parameters may not be, because different choices of contrasts include non-equivalent priors. We have made the allele-symmetric choice because it is insensitive to the choice of reference allele and because it fits more naturally in existing schemes for computing with tree sequences.

To see this, consider the (contrived) case in which we have *n* haploid samples genotyped at a single locus, and every sample has a distinct genotype. If we write *Z*_*k*_ as the effect of genotype *k*, and *Y* for an intercept, then *X*_*k*_, the trait value of sample *k* is

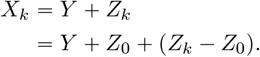

Now, the two models are:

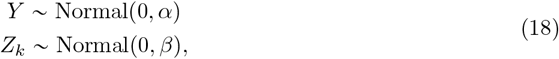

and

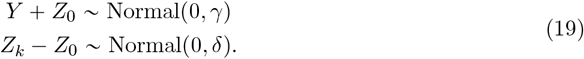

Under model (18), the covariance matrix of *X* is

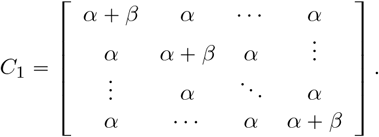

On the other hand, under model (19), the covariance matrix of *X* is

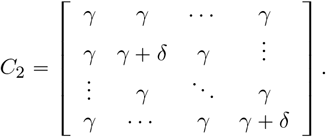

For *n* = 2 and a given *α* and *β*, we can choose *γ* and *δ* so that the two matrices are the same; however, this is not in general possible for *n* > 2, i.e., in the more-than-biallelic case.

One might hope that even though the two models are not equivalent for (*X*_*i*_), they can be made equivalent for the centered values 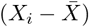. However, one can check that this is not the case: in model (19), the variance of 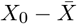 (the ancestral-allele-carrying sample) differs from the other samples, while under model (18) they are the same. To see this, compare *PC*_1_*P* and *PC*_2_*P*, with *P* = *I* − 11^*T*^ /*n*.

#### B Multi-allelic loci in haploids

Following from the methods in the main paper, here we expand the haploid case with two alleles to multiple alleles. The allele of individual *i* at locus *l* is *G*_*i,l*_ ∈ *A* for some alphabet *A*, and each allele *a* at each locus *l* has an additive effect *Z*_*l,a*_. We have this information for *n*_*l*_ loci, *n*_*a*_ distinct alleles, and *n*_*i*_ haploid individuals. (Here we take the alphabet to be the same for all loci, but this is only for convenience, because alleles not present at a locus have no effect.) Recall that the genetic value of individual *i* is

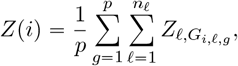

where each allele *a* at each locus ℒ has an independent effect *Z*_ℒ,*a*_, with mean 0 and variance *σ*^2^. However, this might seem not very well defined, since addition of invariant sites affects the result. So, suppose at each locus there is an ancestral allele

We can define the covariance for this case using equation (5). To write this covariance as a sum over alleles, it will be convenient to use the following lemma.

**Lemma 1**. *Let a, b, c, d* ∈ {0, 1}, *and let* [*a* = *b*] = 1 *if a* = *b and* [*a* = *b*] = 0 *otherwise. Then we have:*

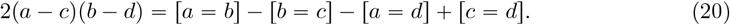

*Proof*. First, notice that both sides are equal to 0 if *a* = *b* = *c* = *d* or if three agree and only one differs. On the right-hand side, this occurs because each of *a, b, c, d* appears in two terms, one positive and one negative. Therefore, if any three agree and differ from a fourth one, then we are left with 1 − 1 − 0 + 0 = 0. Now consider the case of two pairs. If *a* = *b* ≠ *c* = *d*, then both sides are equal to +2. If *a* = *d* ≠ *b* = *c*, then both sides are equal to −2. Finally, if *a* = *c* ≠ *b* = *d*, then both sides are 0.

Now, for each *a* ∈ *A* let 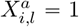 if *G*_*i,l*_ = *a*, and equal 0 otherwise. We have the following identity between individuals *i* and *j*:

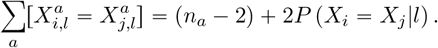

This identity follows from:

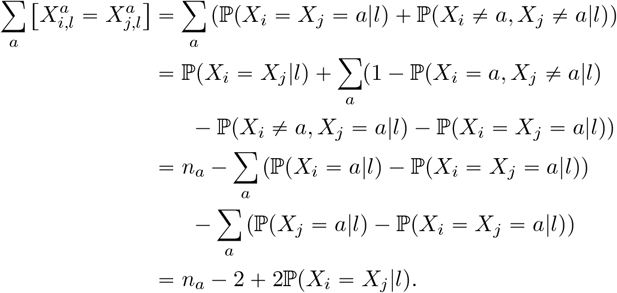

Then, using Lemma 1 we can write (5) as:

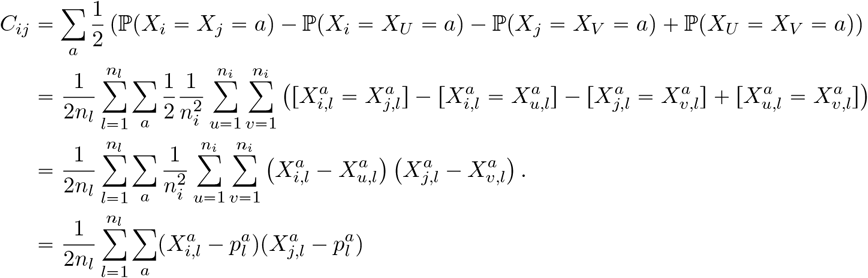

Note that again this expression agrees with equation (2), after dividing the equation by *n*_*l*_.

#### C Proof of equation (5)

As in the main text, *i* and *j* are fixed haploid individuals, while *U* and *V* are uniformly chosen haploid individuals (chosen with replacement, so it may be that *i* = *U* = *V*, for instance). Now, form the random variable (*X*_*i*_, *X*_*j*_, *X*_*U*_, *X*_*V*_) that takes the value (*G*_*i*,ℒ_, *G*_*j*,ℒ_, *G*_*U*,ℒ_, *G*_*V*,ℒ_) with probability 1/ (*n*_*I*_ ^2^*n*_*L*_) for *U, V* = 1, …, *n*_*I*_ and ℒ = 1, …, *n*_*L*_. In other words, we choose the individuals *U* and *V* uniformly at random, with replacement, from the set of *n*_*I*_ individuals, and also choose a locus ℒ uniformly at random from the set of *n*_*L*_ loci; then (*X*_*i*_, *X*_*j*_, *X*_*U*_, *X*_*V*_) is the alleles of those individuals at that locus. In the following, we will repeatedly use the fact that 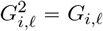 (since *G*_*i*,ℒ_ ∈ {0, 1}). Conditional on locus ℒ, we have:

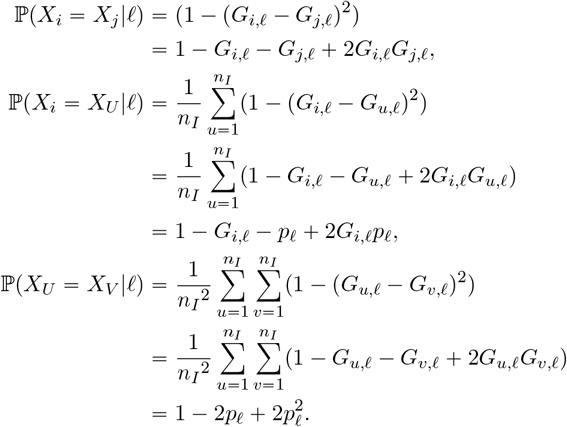

Combining these expressions, we have the following identity:

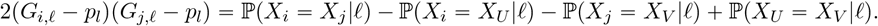

Note the similarity to expression (3). Since ℒ is chosen uniformly at random, it follows that:

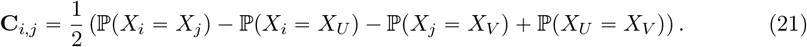

This is expression (5).

**Figure A1:**
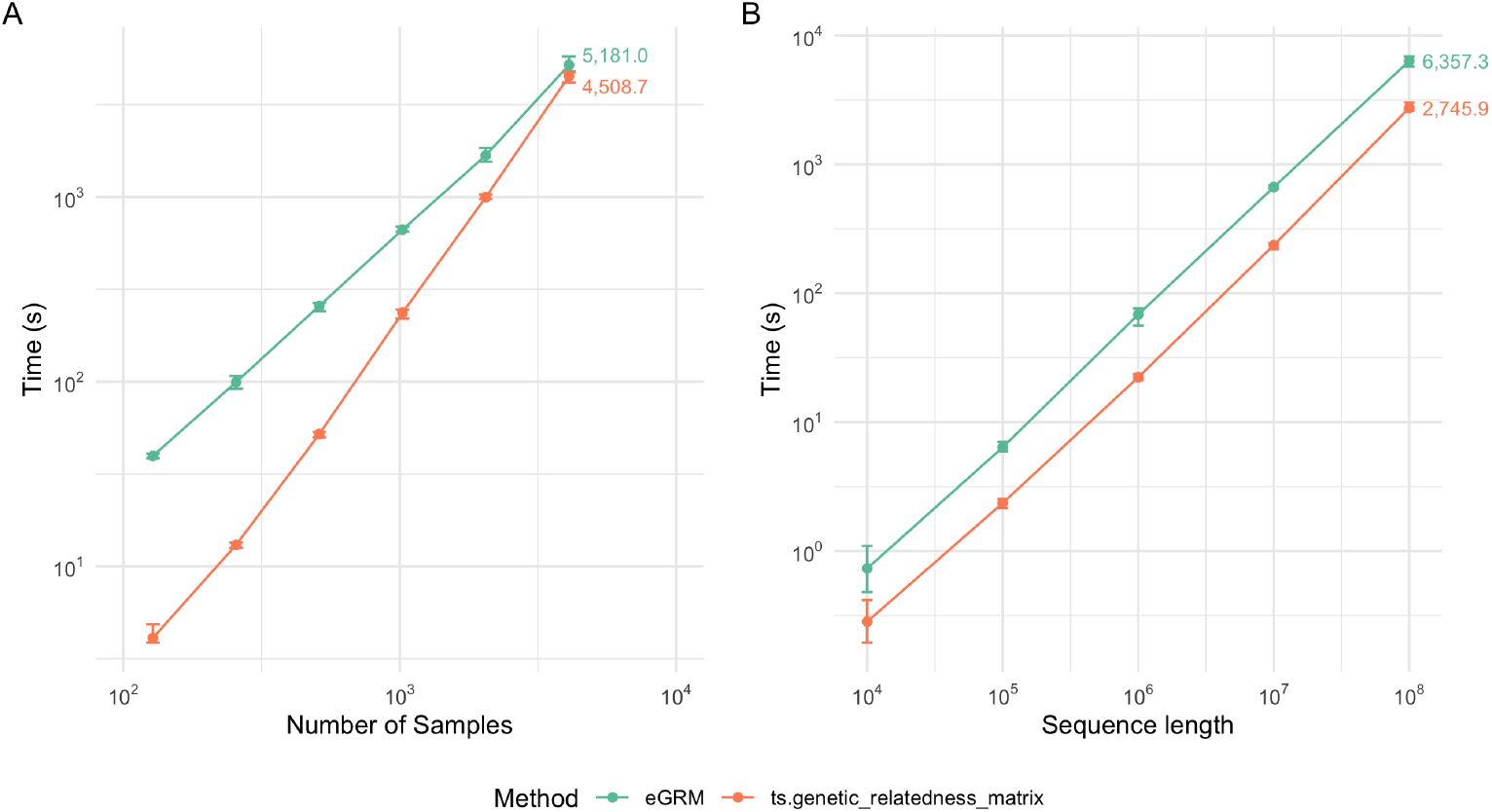
Time efficiency of different implementations of branch GRM computation. Each dot corresponds to the average time taken across ten simulations with different random seeds. Error bars represent the range in time taken across the ten simulations. (A) Branch GRM computation with genome sequence length fixed at 10^7^ and varying the number of samples. (B) Branch GRM computation with number of sample nodes fixed at 2^10^ and varying genome sequence length.

#### D Benchmarking branch GRM computations

#### E Proof of correctness of Algorithm V

Here we prove that when Algorithm V completes, *v*(*s*) is equal to the *s*^th^ entry of **Cw**, as defined in (16). In fact, after each step in the algorithm (i.e., after each addition or removal of an edge), it is true that for *every* node *n*, the sum of everything above that node is equal to the weighted sum of covariances for that node including everything up to that point in the genome. In other words, for every *n*,

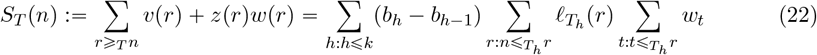

where *T* is the current tree. This statement reduces to our claim that the algorithm is correct because the final tree is an “empty” tree with no edges, so at the end of the algorithm, the left-hand side is simply *v*(*n*) + *z*(*n*). This is again *v*(*n*) because *z*(*n*) is zero due to *x*(*n*) = *b*_*K*_. The right-hand side is equal to equation (16) when *n* is a sample node.

Each time we add an edge with child *c* and parent *p* to the tree (step V2), we add the value of *w*(*n*) to *p* and all nodes above *p* in the tree; when removing edges we subtract(step V1). Since *w*(*n*) is initialized so that each sample *s* carries *w*_*s*_, this ensures that *w* (*n*) = ∑_*s*≤*n*_ *w*_*s*_ at all times [as in Kelleher et al. [2016], Ralph et al. [2020]].

We prove that equation (22) is always true by induction. At the first (empty) tree, this is certainly true, as both sides are equal to zero. We now consider **Step V3**. Tree *T* and the bookkeeping variables *v, w* and *x* are left constant. Advancing the position from *k* to *k* + 1 only changes *z* (*s*) = ℒ_*T*_ (*s*) (*b*_*k*_ − *x* (*s*)) to 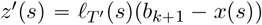. Therefore, the appended value to the left-hand side is

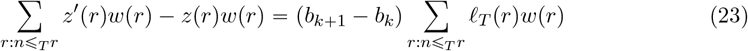

which is also equal to the added value to the right-hand side. Note that this is the only step that changes the value of equation (22).

**Step V1** and **V2** leave the value of both sides unchanged. Without the loss of generality, we prove this for **V2**. Suppose that we added edge (*c, p*) where *c* and *p* are the child and the parent nodes of the edge, respectively. We can divide the nodes of *N* into four categories as (1) the child node *c*, (2) nodes below *c*, (3) the parent node *p* and nodes above *p*, and (4) all other nodes. The insertion changes the values of the intermediate arrays to *v*′, *w*′, *z*′, and *x*′ following **V2**. We denote the new tree resulting from the insertion as *T*′.

Observe that the difference in the left-hand side after **V2** is

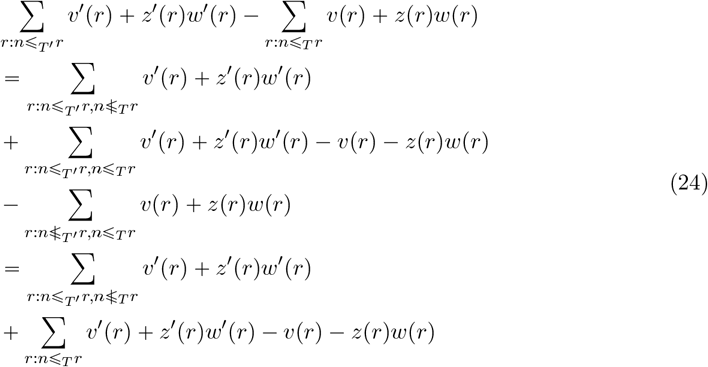

for node *n*. The last line follows from 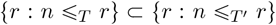 because *T*′ has more edges than *T*. Nodes in each category have distinct values for the former and the latter sum of this equation.

The set 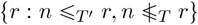 of the first summation is

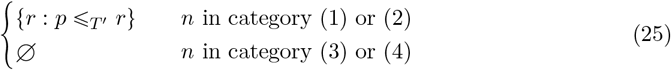

This is because *p* and the nodes above *p* are the nodes that were previously not ancestors in *T*, but became ancestors of *c* and those below *c* after the addition of the new edge. The set is empty for the nodes in the third and the fourth category because their ancestor nodes are unchanged after edge insertion. Therefore, the first summation is

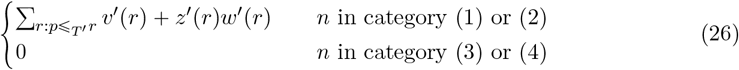

The summand of the second summation *v*′(*r*) + *z*′(*r*) − *v*(*r*) − *z*(*r*) is

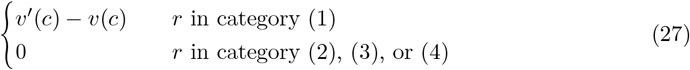

For *r* = *c* (*r* is in the first category), it follows from *z*(*c*) = 0 due to ℒ_*T*_ (*c*) = 0 and *z*′(*c*) = 0 due to *x*′(*c*) = *b*_*k*_. All the bookkeeping values of the second and the fourth category is unchanged by **V2**, so the summand is trivially zero. When *r* = *p* (*r* belongs to the third category), *v*′(*r*) = *v*(*r*) + *z*(*r*)*w*(*r*) and *z*′ (*r*) = 0 by the operations *v*(*r*) + = *z*(*r*)*w*(*r*) and *x*(*r*) = *b*_*k*_ by **V2**. Hence, the second summation is

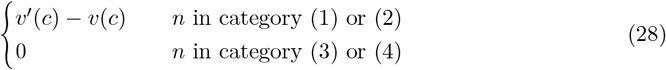

because the set {*r* : *n* ≤_*T*_ *r*} contains *c* if and only if *n* belongs to either the third or the fourth category. An expression for *v*′(*c*) − *v*(*c*) comes from the operation *v*(*c*) − = *v*(*r*) following *v*(*r*) + =*z*(*r*)*w*(*r*) of **V2**

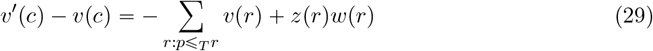

Combining the aforementioned results, we see that equation (24) is

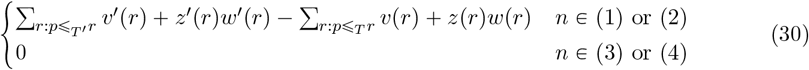

Both cases reduces to zero because the set of nodes ancestral to *p* are the same in *T* and 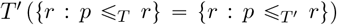, and *v*′ (*r*) = *v*(*r*) + *z*(*r*)*w*(*r*) for these nodes due to **V2** and *z*′(*r*) = 0.

The right-hand side remains the same because the operation changes nothing in its expression: it’s the working sum until the previous local tree that it does not contain any of the components of the current tree that is being modified. This completes the proof.

## References

A. Abdellaoui, L. Yengo, K. J. H. Verweij, and P. M. Visscher. 15 years of GWAS discovery: Realizing the promise. The American Journal of Human Genetics, 110(2):179—-194, 2023. doi: 10.1016/j.ajhg.2022.12.011. URL http://dx.doi.org/10.1016/j.ajhg.2022.12.011.

J. R. Adrion, C. B. Cole, N. Dukler, J. G. Galloway, A. L. Gladstein, G. Gower, C. C. Kyriazis, A. P. Ragsdale, G. Tsambos, F. Baumdicker, J. Carlson, R. A. Cartwright, A. Durvasula, I. Gronau, B. Y. Kim, P. McKenzie, P. W. Messer, E. Noskova, D. O. D. Vecchyo, F. Racimo, T. J. Struck, S. Gravel, R. N. Gutenkunst, K. E. Lohmueller, P. L. Ralph, D. R. Schrider, A. Siepel, J. Kelleher, and A. D. Kern. A community-maintained standard library of population genetic models. eLife, 9, June 2020. doi: 10.7554/elife.54967. URLhttps://doi.org/10.7554%2Felife.54967.

All of Us Research Program Genomics Investigators et al. Genomic data in the All of Us research program. Nature, 627(8003):340, 2024.

L. Anderson-Trocmé, D. Nelson, S. Zabad, A. Diaz-Papkovich, I. Kryukov, N. Baya, M. Touvier, B. Jeffery, C. Dina, H. Vézina, J. Kelleher, and S. Gravel. On the genes, genealogies, and geographies of quebec. Science, 380(6647):849–855, 2023. doi: 10.1126/science.add5300. URL https://www.science.org/doi/10.1126/science.add5300.

S. J. Arnold. Evolutionary Quantitative Genetics Paperback. Oxford University Press, Oxford, UK, 2023.

J. D. Backman, A. H. Li, A. Marcketta, D. Sun, J. Mbatchou, M. D. Kessler, C. Benner, D. Liu, A. E. Locke, S. Balasubramanian, et al. Exome sequencing and analysis of 454,787 UK Biobank participants. Nature, 599(7886):628–634, 2021.

F. Baumdicker, G. Bisschop, D. Goldstein, G. Gower, A. P. Ragsdale, G. Tsambos, S. Zhu, B. Eldon, E. C. Ellerman, J. G. Galloway, A. L. Gladstein, G. Gorjanc, B. Guo, B. Jeffery, W. W. Kretzschumar, K. Lohse, M. Matschiner, D. Nelson, N. S. Pope, C.D. Quinto-Cortés, M. F. Rodrigues, K. Saunack, T. Sellinger, K. Thornton, H. van Kemenade, A. W. Wohns, Y. Wong, S. Gravel, A. D. Kern, J. Koskela, P. L. Ralph, and J. Kelleher. Efficient ancestry and mutation simulation with msprime 1.0. Genetics, 220(3):iyab229, 2022. doi: 10.1093/genetics/iyab229. URL https://doi.org/10.1093/genetics/iyab229.

D. J. Begun, A. K. Holloway, K. Stevens, L. W. Hillier, Y.-P. Poh, M. W. Hahn, P. M. Nista, C. D. Jones, A. D. Kern, C. N. Dewey, L. Pachter, E. Myers, and C. H. Langley. Population genomics: Whole-genome analysis of polymorphism and divergence in Drosophila simulans. PLOS Biology, 5(11):1–26, 2007. doi: 10.1371/journal.pbio.0050310. URL https://doi.org/10.1371/journal.pbio.0050310.

D. Y. C. Brandt, C. D. Huber, C. W. K. Chiang, and D. Ortega-Del Vecchyo. The promise of inferring the past using the Ancestral Recombination Graph (ARG). Genome Biology and Evolution, page evae005, 2024. doi: 10.1093/gbe/evae005. URL https://doi.org/10.1093/gbe/evae005.

S. R. Browning and B. L. Browning. Identity by descent between distant relatives: Detection and applications. Annual Review of Genetics, 46(1):617–633, 2012. doi: 10.1146/annurev-genet-110711-155534. URL https://doi.org/10.1146/annurev-genet-110711-155534.

P. R. Burton, D. G. Clayton, L. R. Cardon, N. Craddock, P. Deloukas, A. Duncanson, D. P. Kwiatkowski, M. I. McCarthy, W. H. Ouwehand, N. J. Samani, J. A. Todd, P. Donnelly, J. C. Barrett, P. R. Burton, D. Davison, P. Donnelly, D. Easton, D. Evans, H.-T. Leung, J. L. Marchini, A. P. Morris, C. C. A. Spencer, M. D. Tobin, L. R. Cardon, D. G. Clayton, A. P. Attwood, J. P. Boorman, B. Cant, U. Everson, J. M. Hussey, J. D. Jolley, A. S. Knight, K. Koch, E. Meech, S. Nutland, C. V. Prowse, H. E. Stevens, N. C. Taylor, G. R. Walters, N. M. Walker, N. A. Watkins, T. Winzer, J. A. Todd, W. H. Ouwehand, R. W. Jones, W. L. McArdle, S. M. Ring, D. P. Strachan, M. Pembrey, G. Breen, D. St Clair, S. Caesar, K. Gordon-Smith, L. Jones, C. Fraser, E. K. Green, D. Grozeva, M. L. Hamshere, P. A. Holmans, I. R. Jones, G. Kirov, V. Moskvina, I. Nikolov, M. C. O’Donovan, M. J. Owen, N. Craddock, D. A. Collier, A. Elkin, A. Farmer, R. Williamson, P. McGuffin, A. H. Young, I. N. Ferrier, S. G. Ball, A. J. Balmforth, J. H. Barrett, D. T. Bishop, M. M. Iles, A. Maqbool, N. Yuldasheva, A. S. Hall, P. S. Braund, P. R. Burton, R. J. Dixon, M. Mangino, S. Stevens, M. D. Tobin, J. R. Thompson, N. J. Samani, F. Bredin, M. Tremelling, M. Parkes, H. Drummond, C. W. Lees, E. R. Nimmo, J. Satsangi, S. A. Fisher, A. Forbes, C. M. Lewis, C. M. Onnie, N. J. Prescott, J. Sanderson, C. G. Mathew, J. Barbour, M. K. Mohiuddin, C. E. Todhunter, J. C. Mansfield, T. Ahmad, F. R. Cummings, D. P. Jewell, J. Webster, M. J. Brown, D. G. Clayton, G. M. Lathrop, J. Connell, A. Dominiczak, N. J. Samani, C. A. B. Marcano, B. Burke, R. Dobson, J. Gungadoo, K. L. Lee, P. B. Munroe, S. J. Newhouse, A. Onipinla, C. Wallace, M. Xue, M. Caulfield, M. Farrall, A. Barton,, T. B. i. R. G. Genomics, I. N. Bruce, H. Donovan, S. Eyre, P. D. Gilbert, S. L. Hider, A. M. Hinks, S. L. John, C. Potter, A. J. Silman, D. P. M. Symmons, W. Thomson, J. Worthington, D. G. Clayton, D. B. Dunger, S. Nutland, H. E. Stevens, N. M. Walker, B. Widmer, J. A. Todd, T. M. Frayling, R. M. Freathy, H. Lango, J. R. B. Perry, B. M. Shields, M. N. Weedon, A. T. Hattersley, G. A. Hitman, M. Walker, K. S. Elliott, C. J. Groves, C. M. Lindgren, N. W. Rayner, N. J. Timpson, E. Zeggini, M. I. McCarthy, M. Newport, G. Sirugo, E. Lyons, F. Vannberg, A. V. S. Hill, L. A. Bradbury, C. Farrar, J. J. Pointon, P. Wordsworth, M. A. Brown, J. A. Franklyn, J. M. Heward, M. J. Simmonds, S. C. L. Gough, S. Seal, B. C. Susceptibility Collaboration, M. R. Stratton, N. Rahman, M. Ban, A. Goris, S. J. Sawcer, A. Compston, D. Conway, M. Jallow, M. Newport, G. Sirugo, K. A. Rockett, D. P. Kwiatkowski, S. J. Bumpstead, A. Chaney, K. Downes, M. J. R. Ghori, R. Gwilliam, S. E. Hunt, M. Inouye, A. Keniry, E. King, R. McGinnis, S. Potter, R. Ravindrarajah, P. Whittaker, C. Widden, D. Withers, P. Deloukas, H.-T. Leung, S. Nutland, H. E. Stevens, N. M. Walker, J. A. Todd, D. Easton, D. G. Clayton, P. R. Burton, M. D. Tobin, J. C. Barrett, D. Evans, A. P. Morris, L. R. Cardon, N. J. Cardin, D. Davison, T. Ferreira, J. Pereira-Gale, I. B. Hallgrimsdóttir, B. N. Howie, J. L. Marchini, C. C. A. Spencer, Z. Su, Y. Y. Teo, D. Vukcevic, P. Donnelly, D. Bentley, M. A. Brown, L. R. Cardon, M. Caulfield, D. G. Clayton, A. Compston, N. Craddock, P. Deloukas, P. Donnelly, M. Farrall, S. C. L. Gough, A. S. Hall, A. T. Hattersley, A. V. S. Hill, D. P. Kwiatkowski, C. G. Mathew, M. I. McCarthy, W. H. Ouwehand, M. Parkes, M. Pembrey, N. Rahman, N. J. Samani, M. R. Stratton, J. A. Todd, and J. Worthington. Genome-wide association study of 14,000 cases of seven common diseases and 3,000 shared controls. Nature, 447(7145):661—-678, 2007. doi: 10.1038/nature05911. URL http://dx.doi.org/10.1038/nature05911.

C. Bycroft, C. Freeman, D. Petkova, G. Band, L. T. Elliott, K. Sharp, A. Motyer, D. Vukcevic, O. Delaneau, J. O’Connell, A. Cortes, S. Welsh, A. Young, M. Effingham, G. McVean, S. Leslie, N. Allen, P. Donnelly, and J. Marchini. The UK Biobank resource with deep phenotyping and genomic data. Nature, 562(7726):203––209, 2018. doi: 10.1038/s41586-018-0579-z. URL https://doi.org/10.1038/s41586-018-0579-z.

M. Caulfield, J. Davies, M. Dennys, L. Elbahy, T. Fowler, S. Hill, T. Hubbard, L. Jostins, N. Maltby, J. Mahon-Pearson, G. McVean, K. Nevin-Ridley, M. Parker, V. Parry, A. Rendon, L. Riley, C. Turnbull, and K. Woods. National Genomic Research Library v5.1, Genomics England, 2017.

A. Cesarani, D. Lourenco, S. Tsuruta, A. Legarra, E. Nicolazzi, P. VanRaden, and I. Misztal. Multibreed genomic evaluation for production traits of dairy cattle in the United States using single-step genomic best linear unbiased predictor. Journal of Dairy Science, 105:5141–5152, 2022. doi: 10.3168/jds.2021-21505. URL https://doi.org/10.3168/jds.2021-21505.

J. T. Chang. Recent common ancestors of all present-day individuals. Advances in Applied Probability, 31(4):1002—-1026, 1999. doi: 10.1239/aap/1029955256. URL https://www.jstor.org/stable/1428340.

B. Charlesworth and D. Charlesworth. Elements of Evolutionary Genetics. Roberts and Company, Greenwoord Village, Colorado, USA, 2010.

C. C. Cockerham. Group inbreeding and coancestry. Genetics, 56(1):89–104, 1967. doi: 10.1093/genetics/56.1.89. URL https://doi.org/10.1093/genetics/56.1.89.

J. B. Cole, C. F. Baes, S. A. Eaglen, T. J. Lawlor, C. Maltecca, M. S. Ortega, and P. M. VanRaden. Management of genetic defects in dairy cattle populations. Journal of Dairy Science, 2025. doi: 10.3168/jds.2024-26035. URL https://doi.org/10.3168/jds.2024-26035.

J. J. Colleau. An indirect approach to the extensive calculation of relationship coefficients. Genetics Selection Evolution, 34(409), 2002. doi: 10.1186/1297-9686-34-4-409. URL https://doi.org/10.1186/1297-9686-34-4-409.

J.-J. Colleau, I. Palhiére, S.T. Rodríguez-Ramilo, and A. Legarra. A fast indirect method to compute functions of genomic relationships concerning genotyped and ungenotyped individuals, for diversity management. Genetics Selection Evolution, 49(87), 2017. doi:10.1186/s12711-017-0363-9. URL https://gsejournal.biomedcentral.com/articles/10.1186/s12711-017-0363-9.

M. B. Cook, S. C. Sanderson, J. E. Deanfield, F. Reddington, A. Roddam, D. J. Hunter, and R. Ali. Our Future Health: a unique global resource for discovery and translational research. Nature Medicine, 2025. doi: 10.1038/s41591-024-03438-0.

C. W. Cotterman. A Calculus for Statistico-genetics. PhD thesis, Ohio State University, Columbus, Ohio, USA, 1940.

J. F. Crow and M. Kimura. An Introduction to Population Genetics Theory. The Blackburn Press, Caldwell, New Jersey, USA, 2009.

D. Cruden. The computation of inbreeding coefficients: for closed populations. Journal of heredity, 40(9):248–251, 1949. doi: 10.1093/oxfordjournals.jhered.a106039. URL https://doi.org/10.1093/oxfordjournals.jhered.a106039.

D. DeHaas, Z. Pan, and X. Wei. Enabling efficient analysis of biobank-scale data with genotype representation graphs. Nature Computational Science, Dec. 2024. ISSN 2662-8457. doi: 10.1038/s43588-024-00739-9. URL https://doi.org/10.1038/s43588-024-00739-9.

Y. Deng, R. Nielsen, and Y. S. Song. Robust and accurate bayesian inference of genomewide genealogies for large samples. bioRxiv, 2024. doi: 10.1101/2024.03.16.585351. URL https://www.biorxiv.org/content/early/2024/03/16/2024.03.16.585351.

A. Durvasula and K. E. Lohmueller. Negative selection on complex traits limits phenotype prediction accuracy between populations. The American Journal of Human Genetics, 108 (4):620—-631, 2021. doi: 10.1016/j.ajhg.2021.02.013. URL http://dx.doi.org/10.1016/j.ajhg.2021.02.013.

D. Edwards. Two molecular measures of relatedness based on haplotype sharing. BMC Bioinformatics, 16(383), 2015. doi: 10.1186/s12859-015-0802-y. URL http://dx.doi.org/10.1186/s12859-015-0802-y.

L. O. Emik and C. E. Terrill. Systematic procedures for calculating inbreeding coefficients. Journal of Heredity, 40(2):51–55, 1949. doi: 10.1093/oxfordjournals.jhered.a105986. URL https://doi.org/10.1093/oxfordjournals.jhered.a105986.

D. S. Falconer and T.F.C. Mackay. Introduction to Quantitative Genetics. Longman, Harlow, UK, 1996.

C. Fan, N. Mancuso, and C.W.K. Chiang. A genealogical estimate of genetic relationships. The American Journal of Human Genetics, 109(5):812–824, 2022. doi: 10.1016/j.ajhg.2022.03.016. URL https://www.sciencedirect.com/science/article/pii/S0002929722001124.

J. Felsenstein. Phylogenies and the comparative method. The American Naturalist, 125(1):pp. 1–15, 1985. ISSN 00030147. URL http://www.jstor.org/stable/2461605.

R. A. Fisher. The correlation between relatives on the supposition of Mendelian inheritance. Earth and Environmental Science Transactions of the Royal Society of Edinburgh, 52(2):399–433, 1919. doi: 10.1017/S0080456800012163. URL https://doi.org/10.1017/S0080456800012163.

L.A. García-Cortés, A. Legarra, C. Chevalet, and M.Á. Toro. Variance and covariance of actual relationships between relatives at one locus. PLOS ONE, 8(2):1–5, 2013. doi: 10.1371/journal.pone.0057003. URL https://doi.org/10.1371/journal.pone.0057003.

M. Grossman and E. J. Eisen. Inbreeding, coancestry, and covariance between relatives for X-chromosomal loci. Journal of Heredity, 80(2):137–142, 1989. doi: 10.1093/oxfordjournals.jhered.a110812. URL https://doi.org/10.1093/oxfordjournals.jhered.a110812.

A. F. Gunnarsson, J. Zhu, B. C. Zhang, Z. Tsangalidou, A. Allmont, and P. F. Palamara. A scalable approach for genome-wide inference of ancestral recombination graphs. bioRxiv, 2024. doi: 10.1101/2024.08.31.610248. URL https://www.biorxiv.org/content/early/2024/09/02/2024.08.31.610248.

N. Halko, P. G. Martinsson, and J. A. Tropp. Finding structure with randomness: Probabilistic algorithms for constructing approximate matrix decompositions. SIAM Review, 53(2):217—-288, 2011. doi: 10.1137/090771806. URL http://dx.doi.org/10.1137/090771806.

B. V. Halldorsson, H. P. Eggertsson, K. H. Moore, H. Hauswedell, O. Eiriksson, M. O. Ulfarsson, G. Palsson, M. T. Hardarson, A. Oddsson, B. O. Jensson, et al. The sequences of 150,119 genomes in the UK Biobank. Nature, 607(7920):732–740, 2022.

K. Harris. Using enormous genealogies to map causal variants in space and time. Nature Genetics, 55:730–731, 2023. doi: 10.1038/s41588-023-01389-9. URL https://doi.org/10.1038/s41588-023-01389-9.

C. R. Henderson. A simple method for computing the inverse of a numerator relationship matrix used in prediction of breeding values. Biometrics, 32(1):69–83, 1976. doi: 10.2307/2529339. URL https://www.jstor.org/stable/2529339.

C. R. Henderson. Applications of Linear Models in Animal Breeding. University of Guelph, Guelph, Ontario, Canada, 1984.

J. Hickey, B. Kinghorn, B. Tier, S. Clark, J. van der Werf, and G. Gorjanc. Genomic evaluations using similarity between haplotypes. Journal of Animal Breeding and Genetics, 130(4):259– 269, 2013. doi: 10.1111/jbg.12020. URL https://onlinelibrary.wiley.com/doi/abs/10.1111/jbg.12020.

W. G. Hill and B. S. Weir. Variation in actual relationship as a consequence of mendelian sampling and linkage. Genetics Research, 93(1):47—-64, 2011. doi: 10.1017/S0016672310000480. URL https://doi.org/10.1017/S0016672310000480.

A. Jacquard. Inbreeding: One word, several meanings. Theoretical population biology, 7 (3):338–363, 1975. doi: 10.1016/0040-5809(75)90024-6. URL https://doi.org/10.1016/0040-5809(75)90024-6.

J. Kelleher and K. Lohse. Coalescent simulation with msprime. In J. Y. Dutheil, editor, Statistical Population Genomics, pages 191–230. Springer US, New York, NY, 2020.

J. Kelleher, A. M. Etheridge, and G. McVean. Efficient coalescent simulation and genealogical analysis for large sample sizes. PLOS Computational Biology, 12(5):e1004842, 2016. doi: 10.1371/journal.pcbi.1004842. URL http://dx.doi.org/10.1371/journal.pcbi.1004842.

J. Kelleher, K. R. Thornton, J. Ashander, and P. L. Ralph. Efficient pedigree recording for fast population genetics simulation. PLOS Computational Biology, 14(11):e1006581, 2018. doi: 10.1371/journal.pcbi.1006581. URL http://dx.doi.org/10.1371/journal.pcbi.1006581.

J. Kelleher, Y. Wong, A. W. Wohns, C. Fadil, P. K. Albers, and G. McVean. Inferring whole-genome histories in large population datasets. Nature Genetics, 51(9):1330– 1338, 2019. doi: 10.1038/s41588-019-0483-y. URL https://www.nature.com/articles/s41588-019-0483-y.

J. Kelleher, B. Jeffery, Y. Wong, P. Ralph, G. Tsambos, S. H. Zhan, K. R. Thornton, D. Goldstein, N. Pope, A. W. Wohns, K. Lohmueller, D. Mbuli-Robertson, G. Gower, H. van Kemenade, B. Zhang, M. F. Rodrigues, C. Ellerman, D. Palmer, C. Weiss, G. Bisschop, J. Guez, S. Karthikeyan, A. Zhang, I. Rebollo, S. Belsare, and A. Kern. tskit: the tree sequence toolkit. Zenodo, 2024. URL 10.5281/zenodo.13941739.

D. E. Knuth. Combinatorial Algorithms, Part 1, volume 4A of The Art of Computer Programming. Addison-Wesley, Upper Saddle River, New Jersey, 2011.

K. Lange and J. S. Sinsheimer. Calculation of genetic identity coefficients. Annals of human genetics, 56(4):339–346, 1992.

C. H. Langley, K. Stevens, C. Cardeno, Y. C. G. Lee, D. R. Schrider, J. E. Pool, S. A. Langley, C. Suarez, R. B. Corbett-Detig, B. Kolaczkowski, S. Fang, P. M. Nista, A. K. Holloway, A. D. Kern, C. N. Dewey, Y. S. Song, M. W. Hahn, and D. J. Begun. Genomic variation in natural populations of Drosophila melanogaster. Genetics, 192(2):533–598, 2012. doi: 10.1534/genetics.112.142018. URL https://doi.org/10.1534/genetics.112.142018.

A. Legarra, O. F. Christensen, Z. G. Vitezica, I. Aguilar, and I. Misztal. Ancestral relationships using metafounders: Finite ancestral populations and across population relationships. Genetics, 200(2):455–468, 2015. doi: 10.1534/genetics.115.177014. URL https://doi.org/10.1534/genetics.115.177014.

R. B. Lehoucq, D. C. Sorensen, and C. Yang. ARPACK users’ guide: solution of large-scale eigenvalue problems with implicitly restarted Arnoldi methods. SIAM, 1998.

A. L. Lewanski, M. C. Grundler, and G. S. Bradburd. The era of the ARG: An introduction to ancestral recombination graphs and their significance in empirical evolutionary genomics. PLOS Genetics, 20(1):1–24, 2024. doi: 10.1371/journal.pgen.1011110. URL https://doi.org/10.1371/journal.pgen.1011110.

V. Link, J. G. Schraiber, C. Fan, B. Dinh, N. Mancuso, C. W. K. Chiang, and M. D. Edge. Tree-based QTL mapping with expected local genetic relatedness matrices. The American Journal of Human Genetics, 110(12):2077–2091, 2023. doi: 10.1016/j.ajhg.2023.10.017. URL https://www.sciencedirect.com/science/article/pii/S0002929723003956.

M. Lynch. Methods for the analysis of comparative data in evolutionary biology. Evolution, 45(5):1065–1080, 1991. doi: 10.1111/j.1558-5646.1991.tb04375.x. URL https://onlinelibrary.wiley.com/doi/abs/10.1111/j.1558-5646.1991.tb04375.x.

M. Lynch and B. Walsh. Genetics and Analysis of Quantitative Traits. Sinauer Associates, Sunderland, Massachusetts, USA, 1998.

I. M. MacLeod, B. J. Hayes, and M. E. Goddard. The effects of demography and long-term selection on the accuracy of genomic prediction with sequence data. Genetics, 198(4):1671—-1684, 2014. doi: 10.1534/genetics.114.168344. URL http://dx.doi.org/10.1534/genetics.114.168344.

I. M. MacLeod, P. J. Bowman, C. J. Vander Jagt, M. Haile-Mariam, K. E. Kemper, A. J. Chamberlain, C. Schrooten, B. J. Hayes, and M. E. Goddard. Exploiting biological priors and sequence variants enhances QTL discovery and genomic prediction of complex traits. BMC Genomics, 17(1), 2016. doi: 10.1186/s12864-016-2443-6. URL http://dx.doi.org/10.1186/s12864-016-2443-6.

G. Malécot. Les mathematiques de l’heredite. Masson and Cie, Paris, France, 1948.

G. Malécot. The mathematics of heredity. W. H. Freeman, San Francisco, California, USA, 1969. Yermanos, Demetrios M. (revision, editing, and translation); https://wellcomecollection.org/works/msfaxgkw.

A. Manichaikul, J. C. Mychaleckyj, S. S. Rich, K. Daly, M. Sale, and W.-M. Chen. Robust relationship inference in genome-wide association studies. Bioinformatics, 26(22):2867—-2873, 2010. doi: 10.1093/bioinformatics/btq559. URL https://dx.doi.org/10.1093/bioinformatics/btq559.

A. R. Martin, C. R. Gignoux, R. K. Walters, G. L. Wojcik, B. M. Neale, S. Gravel, M. J. Daly, C. D. Bustamante, and E. E. Kenny. Human demographic history impacts genetic risk prediction across diverse populations. The American Journal of Human Genetics, 100 (4):635––649, 2017. doi: 10.1016/j.ajhg.2017.03.004. URL http://dx.doi.org/10.1016/j.ajhg.2017.03.004.

T. Mary-Huard and D. J. Balding. Fast and accurate joint inference of coancestry parameters for populations and/or individuals. PLOS Genetics, 19(1):e1010054, 2023. doi: 10.1371/journal.pgen.1010054. URL https://dx.doi.org/10.1371/journal.pgen.1010054.

G. McVean. A genealogical interpretation of principal components analysis. PLOS Genetics, 5(10):1–10, 2009. doi: 10.1371/journal.pgen.1000686. URL https://doi.org/10.1371/journal.pgen.1000686.

T. Meuwissen, B. Hayes, and M. Goddard. Accelerating improvement of livestock with genomic selection. Annual Review of Animal Biosciences, 1:221–237, 2013. doi: 10.1146/annurev-animal-031412-103705. URL https://www.annualreviews.org/content/journals/10.1146/annurev-animal-031412-103705.

T. H. E. Meuwissen, B. J. Hayes, and M. E. Goddard. Prediction of total genetic value using genome-wide dense marker maps. Genetics, 157(4):1819—-1829, 2001. doi: 10.1093/genetics/157.4.1819. URL https://dx.doi.org/10.1093/genetics/157.4.1819.

A. Miles, M. F. Rodrigues, P. Ralph, J. Kelleher, M. Schelker, R. Pisupati, S. Rae, and T. Millar. scikit-allel: Explore and analyse genetic variation. Zenodo, 2024. URL 10.5281/zenodo.10876220.

R. Mrode and I. Pocrnic. Linear Models for the Prediction of the Genetic Merit of Animals. CABI, Wallingford, UK, 2023.

J. Nait Saada, G. Kalantzis, D. Shyr, F. Cooper, M. Robinson, A. Gusev, and P. F. Palamara. Identity-by-descent detection across 487,409 British samples reveals fine scale population structure and ultra-rare variant associations. Nature Communications, 11(1), 2020. doi:10.1038/s41467-020-19588-x. URL http://dx.doi.org/10.1038/s41467-020-19588-x.

R. Nielsen, A. H. Vaughn, and Y. Deng. Inference and applications of ancestral recombination graphs. Nature Reviews Genetics, 2024. doi: 10.1038/s41576-024-00772-4. URL https://doi.org/10.1038/s41576-024-00772-4.

A. Ochoa and J. D. Storey. Estimating FST and kinship for arbitrary population structures. PLOS Genetics, 17(1):e1009241, 2021. doi: 10.1371/journal.pgen.1009241. URL https://dx.doi.org/10.1371/journal.pgen.1009241.

T. Pook, M. Schlather, G. de los Campos, M. Mayer, C. C. Schoen, and H. Simianer. HaploBlocker: Creation of subgroup-specific haplotype blocks and libraries. Genetics, 212(4):1045––1061, 2019. doi: 10.1534/genetics.119.302283. URL http://dx.doi.org/10.1534/genetics.119.302283.

P. Ralph, K. Thornton, and J. Kelleher. Efficiently summarizing relationships in large samples: A general duality between statistics of genealogies and genomes. Genetics, 215(3):779–797, 2020. doi: 10.1534/genetics.120.303253. URL https://doi.org/10.1534/genetics.120.303253.

P. L. Ralph. An empirical approach to demographic inference with genomic data. Theoretical Population Biology, 127:91 – 101, 2019. ISSN 0040-5809. doi: 10.1016/j.tpb.2019.03.005. URL http://www.sciencedirect.com/science/article/pii/S0040580918301667.

M. D. Rasmussen, M. J. Hubisz, I. Gronau, and A. Siepel. Genome-wide inference of ancestral recombination graphs. PLOS Genetics, 10(5):e1004342, 2014.

R. Ros-Freixedes, M. Johnsson, A. Whalen, C.-Y. Chen, B. D. Valente, W. O. Herring, G. Gorjanc, and J. M. Hickey. Genomic prediction with whole-genome sequence data in intensely selected pig lines. Genetics Selection Evolution, 54(1), 2022a. doi: 10.1186/s12711-022-00756-0. URL http://dx.doi.org/10.1186/s12711-022-00756-0.

R. Ros-Freixedes, B. D. Valente, C.-Y. Chen, W. O. Herring, G Gorjanc, J. M. Hickey, and M. Johnsson. Rare and population-specific functional variation across pig lines. Genetics Selection Evolution, 54:39, 2022b. doi: 10.1186/s12711022007328.

B. Schieber and U. Vishkin. On finding lowest common ancestors: Simplification and parallelization. SIAM Journal on Computing, 17(6):1253––1262, 1988. doi: 10.1137/0217079. URL http://dx.doi.org/10.1137/0217079.

J. G. Schraiber, M. D. Edge, and M. Pennell. Unifying approaches from statistical genetics and phylogenetics for mapping phenotypes in structured populations. bioRxiv, 2024. doi: 10.1101/2024.02.10.579721v1. URL https://www.biorxiv.org/content/10.1101/2024.02.10.579721v1.

C. Semple and M. Steel. Phylogenetics. Oxford University Press, 2003.

M. Slatkin. Inbreeding coefficients and coalescence times. Genetical Research, 58(2):167—-175, 1991. doi: 10.1017/S0016672300029827. URL https://doi.org/10.1017/S0016672300029827.

S. P. Smith and F. R. Allaire. Efficient selection rules to increase non-linear merit: application in mate selection. Genetics Selection Evolution, 17(3):387–406, 1985. doi: 10.1186/1297-9686-17-3-387. URL https://doi.org/10.1186/1297-9686-17-3-387.

D. Speed and D. J. Balding. Relatedness in the post-genomic era: is it still useful? Nature Reviews Genetics, 16(1):33–44, 2015. doi: 10.1038/nrg3821. URL https://www.nature.com/articles/nrg3821.

D. Speed, G. Hemani, M. R. Johnson, and D. J. Balding. Improved heritability estimation from genome-wide SNPs. The American Journal of Human Genetics, 91(6):1011—-1021, 2012. doi: 10.1016/j.ajhg.2012.10.010. URL https://dx.doi.org/10.1016/j.ajhg.2012.10.010.

L. Speidel, M. Forest, S. Shi, and S. R. Myers. A method for genome-wide genealogy estimation for thousands of samples. Nature Genetics, 51(9):1321–1329, 2019. doi: 10.1038/s41588-019-0484-x. URL https://dx.doi.org/10.1038/s41588-019-0484-x.

Z. Stark, D. Glazer, O. Hofmann, A. Rendon, C. R. Marshall, G. S. Ginsburg, C. Lunt, N. Allen, M. Effingham, J. Hastings Ward, et al. A call to action to scale up research and clinical genomic data sharing. Nature Reviews Genetics, pages 1–7, 2024.

J. Tang and C. W. K. Chiang. A genealogy-based approach for revealing ancestry-specific structures in admixed populations. bioRxiv, 2025. doi: 10.1101/2025.01.10.632475. URL http://dx.doi.org/10.1101/2025.01.10.632475.

E. A. Thompson. The estimation of pairwise relationships. Annals of Human Genetics, 39(2):173–188, 1975. doi: 10.1111/j.1469-1809.1975.tb00120.x. URL https://doi.org/10.1111/j.1469-1809.1975.tb00120.x.

E. A. Thompson. Identity by descent: Variation in meiosis, across genomes, and in populations. Genetics, 194(2):301–326, 2013. doi: 10.1534/genetics.112.148825. URL https://doi.org/10.1534/genetics.112.148825.

G. Tsambos. Efficient analysis of genetic ancestry in population-sized datasets. PhD thesis, The University of Melbourne, Melbourne, Victoria, Australia, 2022.

C. Turnbull, R. H. Scott, E. Thomas, L. Jones, N. Murugaesu, D. Pretty, Freya Boardmanand Halai, E. Baple, C. Craig, A. Hamblin, S. Henderson, C. Patch, et al. The 100 000 Genomes Project: bringing whole genome sequencing to the NHS. BMJ, 361:k1687, 2018a.

C. Turnbull, R. H. Scott, E. Thomas, L. Jones, N. Murugaesu, F. B. Pretty, D. Halai, E. Baple, C. Craig, A. Hamblin, S. Henderson, C. Patch, A. O’Neill, A. Devereau, K. Smith, A. R. Martin, A. Sosinsky, E. M. McDonagh, R. Sultana, M. Mueller, D. Smedley, A. Toms, L. Dinh, T. Fowler, M. Bale, T. Hubbard, A. Rendon, S. Hill, M. J. Caulfield, and 100 000 Genomes Project. The 100 000 Genomes Project: bringing whole genome sequencing to the NHS. BMJ, 361:k1687, 2018b. doi: 10.1136/bmj.k1687. URL https://www.bmj.com/content/361/bmj.k1687.

UK Biobank Whole-Genome Sequencing Consortium, S. Li, K. J. Carss, B. V. Halldorsson, and A. Cortes. Whole-genome sequencing of half-a-million UK Biobank participants. medRxiv, pages 2023–12, 2023.

P. M. VanRaden. Efficient methods to compute genomic predictions. Journal of Dairy Science, 91(11):4414–4423, 2008. doi: 10.3168/jds.2007-0980. URL https://linkinghub.elsevier.com/retrieve/pii/S0022030208709901.

H. Vézina and J.-S. Bournival. An overview of the BALSAC population database. Past developments, current state and future prospects. Historical Life Course Studies, 2020.

P. Virtanen, R. Gommers, T. E. Oliphant, M. Haberland, T. Reddy, D. Cournapeau, E. Burovski, P. Peterson, W. Weckesser, J. Bright, S. J. van der Walt, M. Brett, J. Wilson, K. J. Millman, N. Mayorov, A. R. J. Nelson, E. Jones, R. Kern, E. Larson, C. J. Carey, I?. Polat, Y. Feng, E. W. Moore, J. VanderPlas, D. Laxalde, J. Perktold, R. Cimrman, I. Henriksen, E. A. Quintero, C. R. Harris, A. M. Archibald, A. H. Ribeiro, F. Pedregosa, P. van Mulbregt, A. Vijaykumar, A. P. Bardelli, A. Rothberg, A. Hilboll, A. Klöckner, A. Scopatz, A. Lee, A. Rokem, C. N. Woods, C. Fulton, C. Masson, C. Häggström, C. Fitzgerald, D. A. Nicholson, D. R. Hagen, D. V. Pasechnik, E. Olivetti, E. Martin, E. Wieser, F. Silva, F. Lenders, F. Wilhelm, G. Young, G. A. Price, G.-L. Ingold, G. E. Allen, G. R. Lee, H. Audren, I. Probst, J. P. Dietrich, J. Silterra, J. T. Webber, J. Slavič, J. Nothman, J. Buchner, J. Kulick, J. L. Schönberger, J. V. de Miranda Cardoso, J. Reimer, J. Harrington, J. L. C. Rodríguez, J. Nunez-Iglesias, J. Kuczynski, K. Tritz, M. Thoma, M. Newville, M. Kümmerer, M. Bolingbroke, M. Tartre, M. Pak, N. J. Smith, N. Nowaczyk, N. Shebanov, O. Pavlyk, P. A. Brodtkorb, P. Lee, R. T. McGibbon, R. Feldbauer, S. Lewis, S. Tygier, S. Sievert, S. Vigna, S. Peterson, S. More, T. Pudlik, T. Oshima, T. J. Pingel, T. P. Robitaille, T. Spura, T. R. Jones, T. Cera, T. Leslie, T. Zito, T. Krauss, U. Upadhyay, Y. O. Halchenko, and Y. Vázquez-Baeza. SciPy 1.0: Fundamental algorithms for scientific computing in Python. Nature Methods, 17(3):261–272, 2020. doi: 10.1038/s41592-019-0686-2. URL http://dx.doi.org/10.1038/s41592-019-0686-2.

P. M. Visscher, S. E. Medland, M. A. R. Ferreira, K. I. Morley, G. Zhu, B. K. Cornes, G. W. Montgomery, and N. G. Martin. Assumption-free estimation of heritability from genomewide identity-by-descent sharing between full siblings. PLOS Genetics, 2(3):1–10, 2006. doi: 10.1371/journal.pgen.0020041. URL https://doi.org/10.1371/journal.pgen.0020041.

B. Walsh and M. Lynch. Evolution and Selection of Quantitative Traits. Oxford University Press, Oxford, UK, 2018.

Y. Wang, K. Tsuo, M. Kanai, B. M. Neale, and A. R. Martin. Challenges and opportunities for developing more generalizable polygenic risk scores. Annual Review of Biomedical Data Science, 5(1):293–320, 2022. doi: 10.1146/annurev-biodatasci-111721-074830. URL http://dx.doi.org/10.1146/annurev-biodatasci-111721-074830.

B. S. Weir and J. Goudet. A unified characterization of population structure and relatedness. Genetics, 206(4):2085–2103, 2017. doi: 10.1534/genetics.116.198424. URL https://academic.oup.com/genetics/article/206/4/2085/6072590.

B. S. Weir and J. Goudet. How to estimate kinship. Molecular Evolutionary Ecology, 27(20):4121–4135, 2018. doi: 10.1111/mec.14833. URL https://doi.org/10.1111/mec.14833.

B. S. Weir, A. D. Anderson, and A. B. Hepler. Genetic relatedness analysis: modern data and new challenges. Nature Reviews Genetics, 7(10):771––780, 2006. doi: 10.1038/nrg1960. URL https://dx.doi.org/10.1038/nrg1960.

O. Weissbrod, F. Hormozdiari, C. Benner, R. Cui, J. Ulirsch, S. Gazal, A. P. Schoech, B. van de Geijn, Y. Reshef, C. Márquez-Luna, L. O’Connor, M. Pirinen, H. K. Finucane, and A. L. Price. Functionally informed fine-mapping and polygenic localization of complex trait heritability. Nature Genetics, 52(12), Dec. 2020. ISSN 1546-1718. doi: 10.1038/s41588-020-00735-5.

O. Weissbrod, M. Kanai, H. Shi, S. Gazal, W. Peyrot, A. Khera, Y. Okada, K. Y. Y. T. Y. Y. K. A. M. Y. F. M. M. Kamat, K. Matsuda, Y. Yamanashi, Y. Furukawa, T. Morisaki, Y. Murakami, Y. Kamatani, K. Muto, A. Nagai, W. Obara, K. Yamaji, K. Takahashi, S. Asai, Y. Takahashi, T. Suzuki, N. Sinozaki, H. Yamaguchi, S. Minami, S. Murayama, K. Yoshimori, S. Nagayama, D. Obata, M. Higashiyama, A. Masumoto, Y. Koretsune, A. R. Martin, H. Finucane, and A. Price. Leveraging fine-mapping and multi-population training data to improve cross-population polygenic risk scores. Nature Genetics, 54, 2022. doi:10.1038/s41588-022-01036-9.

Y. Wong, A. Ignatieva, J. Koskela, G. Gorjanc, A. W. Wohns, and J. Kelleher. A general and efficient representation of ancestral recombination graphs. Genetics, 228(1):iyae100, 07 2024. doi: 10.1093/genetics/iyae100.

S. Wright. Coefficients of inbreeding and relationship. The American Naturalist, 56(645):330– 338, 1922. doi: 10.1086/279872. URL https://doi.org/10.1086/279872.

S. Wright. The interpretation of population structure by F-statistics with special regard to systems of mating. Evolution, 19(3):395–420, 1965. doi: 10.2307/2406450. URL https://www.jstor.org/stable/2406450.

S. Yair and G. Coop. Population differentiation of polygenic score predictions under stabilizing selection. Philosophical Transactions of the Royal Society B: Biological Sciences, 377(1852), 2022. doi: 10.1098/rstb.2020.0416. URL http://dx.doi.org/10.1098/rstb.2020.0416.

J. Yang, B. Benyamin, B. P. McEvoy, S. Gordon, A. K. Henders, D. R. Nyholt, P. A. Madden, A. C. Heath, N. G. Martin, G. W. Montgomery, M. E. Goddard, and P. M. Visscher. Common SNPs explain a large proportion of the heritability for human height. Nature Genetics, 42 (7):565–569, 2010. doi: 10.1038/ng.608. URL https://doi.org/10.1038/ng.608.

I. Young. Discovering missing heritability in whole-genome sequencing data. Nature Genetics, 54(3):224––226, 2022. doi: 10.1038/s41588-022-01012-3. URL http://dx.doi.org/10.1038/s41588-022-01012-3.

C. Zhang, A. Biddanda, A. F. Gunnarsson, F. Cooper, and P. F. Palamara. Biobank-scale inference of ancestral recombination graphs enables genealogical analysis of complex traits. Nature Genetics, 55(5):768–776, 2023. doi: 10.1038/s41588-023-01379-x. URL https://doi.org/10.1038/s41588-023-01379-x.

J. Zhu, G. Kalantzis, A. Pazokitoroudi, Á. F. Gunnarsson, H. Loya, H. Chen, S. Sankararaman, and P. F. Palamara. Fast variance component analysis using large-scale ancestral recombination graphs. bioRxiv, 2024. doi: 10.1101/2024.08.31.610262. URL https://www.biorxiv.org/content/early/2024/08/31/2024.08.31.610262.

